# Comparing Kinetic versus Stoichiometric Priorities in Hybrid Models of CHO Metabolism

**DOI:** 10.1101/2025.10.28.685134

**Authors:** Pratik Khare, Nelson Ndahiro, Stephanie Klaubert, Edward Ma, Tom Bertalan, Yannis Kevrekidis, Sarah W. Harcum, Michael J. Betenbaugh

## Abstract

Understanding Chinese hamster ovary (CHO) cell metabolism through mathematical models is essential for optimizing culture media and biomanufacturing processes.

Current mechanistic models rely primarily on either flux balance analysis (FBA), estimating intracellular fluxes while assuming steady state, or kinetic modeling, capturing dynamic behavior but typically for a limited number of reactions. Dynamic FBA (dFBA) integrates both approaches in a hybrid framework, but challenges remain in integrating the two formats to describe bioprocesses. In this study, we first enhanced an existing dynamic CHO-metabolism model by incorporating ^13^C-labeled data to refine kinetic expressions and stoichiometric constraints of amino acid pathways, including the asparagine-aspartate network and serine biosynthesis. We next evaluated the impact of prioritizing either stoichiometry, through the pseudo steady state assumption (PSSA), or the kinetic expressions of fluxes. Comparing error and predictive performance for both models for two industrially relevant fed-batch CHO culture conditions involving varying initial concentrations of nutrients and three feed streams, demonstrated that the kinetic-oriented model (KOM) yielded superior predictions for viable cell density (VCD), antibody production, and a range of amino acids and metabolites compared to the stoichiometric oriented model (SOM). Indeed, the KOM was able to predict production-to-consumption shifts of lactate and alanine, fluctuating levels of ammonia based on reversible kinetic expressions, and amino acids like asparagine and the serine-glycine pool. The KOM also provided better predictions for a third case including lactate-supplemented (LS) feed; however, slight parameter adjustments helped to improve model fidelity, likely due to the impact of high lactate on kinetic expressions of antibody (directly) and VCD (indirectly). In summary, our findings demonstrate that hybrid models emphasizing empirical kinetics over strict pseudo-steady-state constraints capture biologically realistic dynamics such as transient shifts for key metabolites like lactate, alanine, and ammonia, and also produce parameters useful across varying conditions, making them a practical and powerful tool for characterizing CHO cell culture performance in the future.

## 1. Introduction

Biopharmaceutical processes account for an important and increasing share of the drug production market^1^. Indeed, biomanufacturing represents nearly 2% of the total US GDP and its fraction is growing at an annual rate of 11%. Monoclonal antibodies, specifically, have become the predominant class of new drugs developed in recent years with the global therapeutic monoclonal antibody market valued at approximately US$115.2 billion in 2018 and predicted to grow to $300 billion by 2025^2,3^. One of the most important tools for the production of these monoclonal antibodies is the Chinese hamster ovary (CHO) cell line. Some of the key reasons why the CHO cell platform accounts for a majority of biopharmaceutical production are high protein titers, robustness in suspension culture across different cell culture environments and process conditions and ease of scalability^4^. With the extensive use of CHO cells in large scale biopharmaceutical production, the need to characterize and optimize cell performance has become a growing area of focus. Computational modeling is one increasingly important approach that biotechnologists are undertaking to improve our understanding of cell culture processes and ultimately optimize the performance of these cell cultures. The key advantage of this technique is that it enables us to get predictions for parameters for certain cell culture conditions while potentially reducing the number of costly and time-consuming experiments. We can then apply these models to predict performance for different conditions as we seek to optimize bio-manufacturing processes.

While there are multiple types of models for cell performance, models that simulate cell metabolism have broadly been used to characterize cell line nutrient consumption, growth, and generation of products such as antibodies as well as metabolic byproducts^5,6^. These computational models are often mechanistic models, in which detailed information about cellular kinetics and metabolism is incorporated, or purely data-driven, or black-box, in which only empirical observations are used to build the data-driven model. While these data driven models allow us to use vast breadths of data collected during experiments, they yield very limited insights into the actual mechanism of the cell culture. Mechanistic models, on the other hand, afford insights into the behavior of our cell culture system under different growth conditions that can be useful in understanding and optimizing these systems. By and large, there are 2 types of mechanistic models that are used to predict cell culture behavior-kinetic models and stoichiometric models^7–9^. Individually, these have several advantages and disadvantages. Stoichiometric models rely on either small-scale or large-scale intracellular metabolic networks. The fluxes are calculated subject to bounds such extracellular uptake/production rates and the reaction stoichiometry, reversibility, flux bounds etc., as additional information. These systems are often solved using a flux balance analysis approach where a pseudo-steady state assumption is employed for the intracellular species and an objective function, often growth or ATP production, is either maximized or minimized^10^. Kinetic models on the other hand, are comprised of ordinary differential equations pertaining to the variables (fluxes or concentrations) of interest. A major distinguishing factor for these types of models is their ability to factor in temporal information instead of solely relying on a steady state^6^. However, given the vast amount of information needed to run these models, these are often limited to smaller sized networks^7,11^

Therefore, a hybrid model combining certain aspects of the kinetic models and stoichiometric models could be viewed as a viable intermediate approach. Such models are often referred to as dynamic flux balance analysis (dFBA) models^12,13^. dFBA models have been used when kinetic information is available and reliable, and where classical FBA models cannot predict with accuracy. These models are advantageous in that it not only predicts the flux distribution at a particular instant of time but can account for dynamic changes between steady states using kinetic expressions of reactions and metabolite concentration dependencies. Indeed, Mahadevan et al. demonstrated the application of dFBA models for analyzing diauxic growth of E. coli^14^. Therefore, in the current study, we have modified and implemented dFBA-like models to analyze the growth, metabolite production and consumption profiles of mammalian (CHO) cells grown in a fed-batch system, under different initial and feeding conditions.

The dFBA model consists of a concise reaction network consisting of key cytosolic and mitochondrial reactions and kinetic expressions of key intracellular reactions, adapted and enhanced from a previous effort^12^. The model tracks metabolite concentrations in the extracellular space and calculates the flux distribution for the entire reaction network iteratively, from start of culture until end of growth phase. The kinetic parameters involved in the reaction expressions are estimated by minimizing the difference between the predicted metabolite concentrations and the measured concentrations. In the sections below, we describe the key components of the model, including the use of ^13^C-labeled data to inform our model, development of the model for various feeding schemes, application of the model to predict experimental conditions for model case studies and comparing the kinetic-oriented dFBA model with the stoichiometric-oriented dFBA model.

## 2. Development of the dynamic flux balance analysis (dFBA) model

Our dFBA model, originally adapted from Nolan et al.^12^, includes 35 reactions and 24 metabolites and encompasses glycolysis, the TCA cycle and multiple non-essential amino acid pathways **(Figure 1)**.

**Figure 1.**
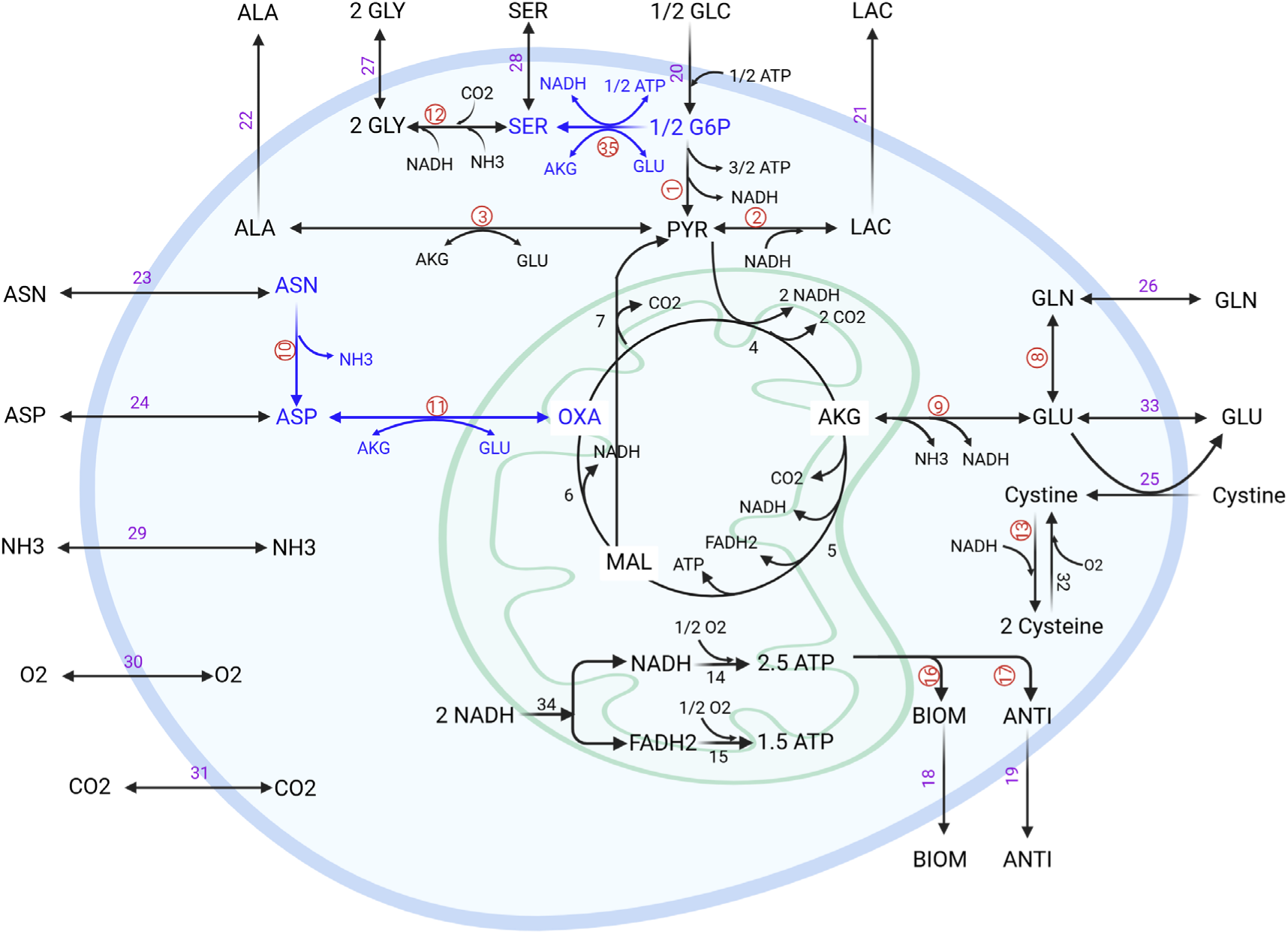
Schematic of the CHO cell metabolic network. The diagram illustrates both intracellular and exchange fluxes. Red-circled reactions represent intracellular reactions modeled using kinetic rate equations. Black reaction numbers denote intracellular reactions modeled with flux balance analysis (FBA), including mitochondrial pathways. Dark blue arrows indicate reactions refined with experimental data from ^13^C-labeled CHO fed-batch cultures. Purple reaction numbers represent exchange fluxes, such as nutrient uptake and metabolite secretion.

### 2.1. Kinetic parameters

Kinetic expressions were constructed for 14 of the 35 reactions, 12 of which were intracellular enzymatic reactions and the remaining 2 were the exchange fluxes for biomass and antibody. Briefly, all the kinetically defined intracellular reactions were modeled based on the Michaelis-Menten format of reaction kinetics. The kinetic expressions were modified from previous work based on additional data collected in our laboratory or from literature. Several modifications were made to the original kinetic expressions and are summarized as follows:

Temperature dependent constants used to scale the maximum forward reaction rate, have been eliminated from the kinetic expressions, as have the inhibition exponential constants, since our dataset of the underlying experimental conditions did not rely on a temperature shift in the operating conditions. Secondly, the redox variable, which was used to account for the redox state of the cell in the reaction kinetics was removed owing to the abrupt discontinuities observed in the reaction expressions, and subsequently, the reaction fluxes. Instead, reactions were modeled solely based on the extracellular metabolites with consumption and production rates being dependent on the concentrations of the associated extracellular metabolites. The kinetic expressions for the 14 reactions are listed in **Supplementary Table 1**, and the complete list of reactions for the modified model is displayed in **Supplementary Table 2.** These two changes also had the benefit of streamlining the parameter estimation as will be discussed in more detail below in **section 2.3**. Removing these temperature-dependent components facilitated parameter estimation, and removing the R term and its associated rate of reactions made the parameter estimation a less complex optimization problem. This made the model more straightforward to use without the need for adjustments requiring excessive computing architecture. Mass balances on all the dynamically tracked species were modeled as forward Euler ODEs. In addition to the various metabolites (glucose, lactate, amino acids) and biomass and antibody, dead cell density was also modeled using an ODE and the associated parameter (death rate) was modeled as a kinetic parameter estimated by the model **(Supplementary Table 3)**.

### 2.2. Modification of reactions via ^13^C-labeled data

Data sets from isotopic carbon (^13^C) labeling experiments were leveraged to modify the model to be more reflective of CHO metabolism. Data collected was used to add pathways that were missing or incomplete in the current model. Specific pathways added/modified to the model are highlighted in dark blue in **Figure 1**. All amino acids were tracked in these ^13^C-labeled datasets, and the behavior of the ones relevant to our model was incorporated. Changes in stoichiometry as well as to the kinetic expressions were implemented resulting in a modified model as described in **sections 3.1.1 and 3.1.2**. The final model consisted of 45 kinetic parameters estimated based on training (control) datasets, further discussed in **section 3.3**.

### 2.3. Implementation of parameter estimation function

In addition to the model modifications, we also implemented additional methods to estimate parameters. To estimate the kinetic parameters, the error between simulated and measured concentrations of dynamically tracked species from the training run was calculated and minimized using the nonlinear optimizer *fmincon*^15^. The error was evaluated using both plain squared error and Symmetric Mean Absolute Percentage Error (SMAPE), with the latter yielding better parameter estimates and therefore selected for final optimization.

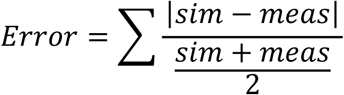

Parameters were estimated in log_10_ space, meaning the optimizer directly adjusted the logarithm of each parameter rather than its raw value. This allowed us to apply simple uniform bounds of [−1,1] in log space, which correspond to multiplicative bounds of [0.1,10] on the original parameters relative to their nominal values. Optimizing in this space ensures a consistent search range across parameters of different magnitudes and improves numerical stability.

Further, we compared the performance of various versions of the model which include:

1. Original implementation of the model
2. Model predictions following improvements in the kinetic expressions and parameters (including elimination of the redox variable).
3. Mechanistic modifications, including changes to the stoichiometric network and kinetic expressions following incorporation of the ^13^C-labeled correlations
4. Final iteration of the model with the improved parameter estimation functions.

Specifically, we compared the overall model error at each time point to that of the final model **(Figure 2A)**. The final time-point error was lowest for the model incorporating all improvements, including optimized parameter estimation (green line), compared to the original model. Furthermore, when comparing the total error, calculated as the sum of errors for each metabolite across all time points (**Figure 2B**), we observed a general decrease in error across iterative versions of the model. The only exception was the version in which the kinetic expressions and parameters (redox variable) were modified (red bar).

**Figure 2.**
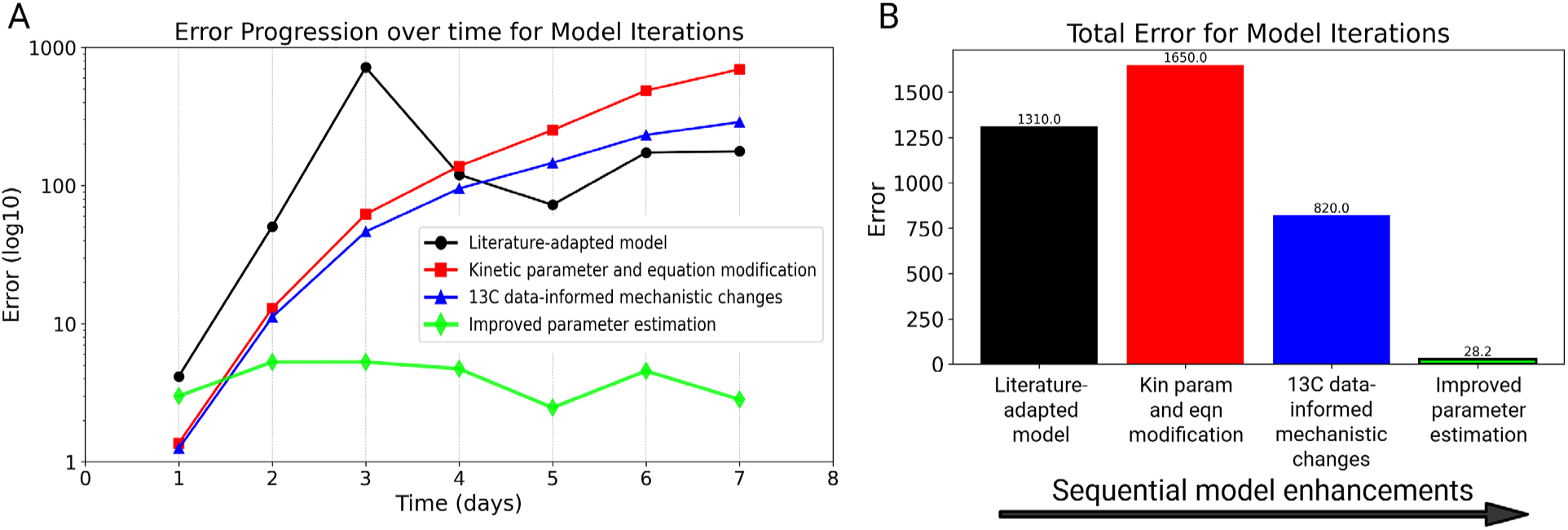
Evolution of model prediction error with successive model refinements. **(A)** Time-course of prediction errors during the 7-day simulation for different model variants. All model iterations, except the improved parameter estimation model iteration, show a generally increasing error trend over time. **(B)** Total prediction error, quantified as the sum of error across all time-points. The improved model demonstrates a substantial reduction in total error, confirming enhanced predictive performance due to model refinement.

The slight increase in prediction error was because the dependency of the kinetic expressions for lactate and alanine production, on the redox variable, were removed.

These expressions had previously modeled the lactate shift by switching between kinetic formulations based on the value of the redox variable. This dependency introduced complexity as flux values became highly sensitive to an arbitrary cutoff. This caused significant fluctuations in predictions between consecutive time points.

Removing the redox variable made the lactate consumption equation entirely dependent on extracellular metabolite concentrations and intracellular fluxes. This allowed the model to operate using a consistent set of equations throughout the simulation. It eliminated abrupt shifts in system behavior that previously resulted from equation switching. As a result, parameter estimation became more efficient because the optimizer only had to fit one set of equations. This simplification also allowed for faster computation and easier implementation of additional features, such as the incorporation of ^13^C data. Although this version showed a slight increase in error, the benefits of simplification and improved parameter estimation outweighed the drawbacks. The error was reduced in later iterations through further optimization. Overall, the iterative versions of the model enhanced its performance, thus validating various mechanistic changes and parameter estimation modifications made to the original model.

### 2.4. Experimental Methods and Materials

The “control case” experiment consisted of a given initial concentration/quantity of each of the dynamically tracked species and three feed streams into the bioreactor labeled as “Feed 7A”, “Feed 7B” (Cell Boost 7a/7b-Cytiva) and “Bolus Glucose Feed” **(Figure S1)**. The initial conditions and concentrations of metabolites included in the model are listed in **Supplementary Tables 4 and 5**. Although the feeds contain additional components, only those relevant to the modeled metabolic network are presented. The experiment was conducted with feed beginning from day 3, i.e., the third day after inoculation. The amount of Feed 7A and 7B differed each day and was fed according to a stepped-pyramid pattern, with a set volume percentage of each of these feeds with respect to the current working volume for the bioreactor **(Figure S1B).** As seen, the feeds were introduced incrementally in a stepwise manner, with increasing percentages of volume fed every few days. Feed 7B is fed at a much (ten times) lower volume % than Feed 7A. The volume of the bolus glucose feed depends on the set-point of glucose **(Figure S1C)**. The volume of bolus glucose thus fed is calculated as:

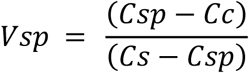

Where, Vsp = Volume of stock needed to be added,

Cs = Concentration of glucose stock

Cc = Concentration of glucose in culture media

Csp = Glucose set point

For the lactate-supplemented case, bolus additions of 150 g/L sodium lactate were added in 10 mM increments at 12, 24, and 36 hours^16^, resulting in a total added concentration of 30 mM.

Cells were precultured in shake flasks and subsequently transferred to an Ambr250 vessel. The ambr250 HT bioreactors (Sartorius Stedim, Göttingen, Germany) were equipped with two pitched blade impellers and an open pipe sparger (vessel part number: 001–5G25). Bioreactors were inoculated to a target VCD of 0.4×10^6^ cells/mL into 210 mL media. Temperature was controlled to 36.5°C. DO was controlled to 50% air saturation using a 6-level proportional-integral-derivative (PID) control algorithm^17^. The control settings for DO are listed in **Supplementary Table 6**. For the control and lactate stressed cultures, samples were taken daily for offline VCD, viability, glucose, glutamate, glutamine, lactate, ammonia, dissolved oxygen (pO_2_), dissolved CO_2_ (pCO_2_), and pH beginning on day 0.

VCD and viability were measured using a Vi-Cell XR cell viability analyzer (Beckman Coulter, Brea, CA). Extracellular glucose, glutamine, glutamate, ammonia, lactate, and IgG concentrations were measured using a Cedex Bio Analyzer (Roche Diagnostics, Mannheim, Germany). Samples for amino acid analysis were centrifuged at 10,000 x g for 10 minutes at 4°C. The supernatant was aliquoted and frozen at-20°C until analysis occurred. Amino acid concentrations were measured using the REBEL cell culture media analyzer (908 Devices, Boston, MA). End of culture harvest samples were taken for glycosylation analysis, which was conducted as described in Synoground et al., 2021^18^. In brief, a Protein A agarose resin was used to obtain the IgG_1_ from the culture broth.

## 3. Results and Discussion

### 3.1. Use of labeled ^13^C data to inform model network

Reaction stoichiometries were revised based on literature as well as ^13^C-isotope-labeled (intracellular and extracellular) metabolite tracking data^19^. Accordingly, the reaction expressions were changed to match the revised stoichiometry. Modified reactions (equations 10, 11) and a new reaction (equation 35, in **Figure 1 and Supplementary Table 1**) were also added to the model with the kinetic expressions devised based on the stoichiometry of those reactions (described in sections below).

Below, we describe how the labeled datasets were used to inform our model.

#### 3.1.1. Asparagine and Aspartate reaction network

Asparagine is a major source of aspartate generation via the deamidation reaction. It has been shown that both asparagine and aspartate are key contributors to the TCA cycle in CHO cells, especially in cases of low glutamate and glutamine supplementation^20–22^. Thus, modeling these reactions appropriately in our reaction network is key to understanding the dynamic flux distribution of both TCA cycle as well as other pathways involving aspartate and asparagine such as protein synthesis and biomass generation. We thus looked at data from ^13^C-labeled asparagine and aspartate experiments performed on CHO-K1 cells. Specifically, we observed the enrichment of aspartate from feeding labeled asparagine to CHO-K1 cells and vice-versa (**Figure 3A**, orange-highlighted trapezoids). While aspartate generation from asparagine was seen, asparagine was not generated from labeled aspartate. Thus, the asparagine deamidation reaction was seen to be either irreversible or proceeding backwards at a very slow rate, resulting in the removal in the reversibility of the reaction in our model and a change in the stoichiometry of the related metabolites (dark blue highlighted pathway in **Figure 3C**).

**Figure 3.**
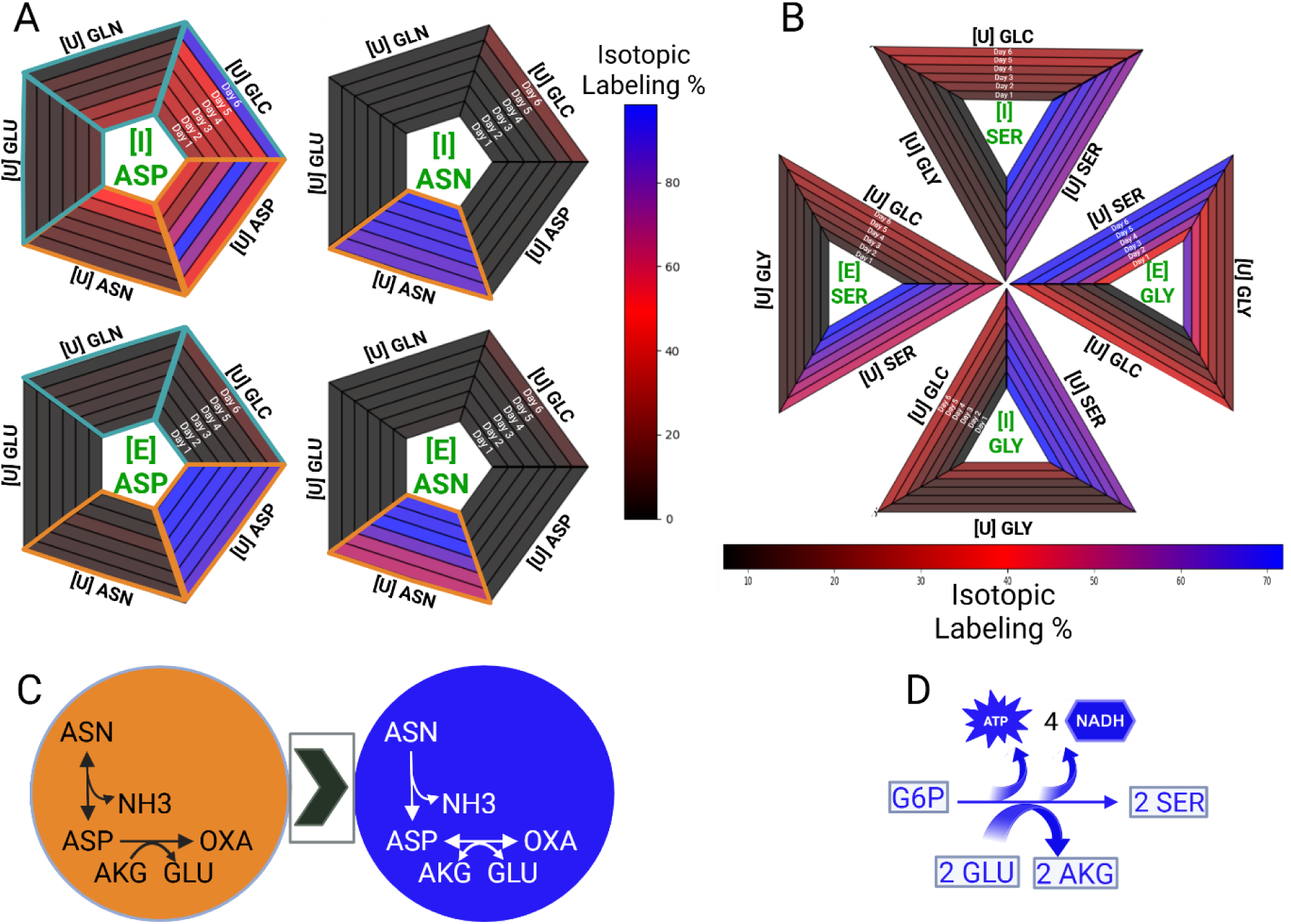
Model refinements based on enrichment of intra-and extracellular metabolites from ^13^C-labeled nutrients in CHO-K1 cell cultures. **(A)** Isotopic enrichment profiles of intracellular [I] and extracellular [E] aspartate and asparagine over a 6-day period following uniform labeling [U] with amino acids (glutamate, glutamine, aspartate, asparagine) and glucose. Teal-highlighted trapezoids indicate datasets that informed modifications to the mechanistic model, specifically the aspartate transamination reaction. Orange-highlighted trapezoids indicate datasets used to refine the asparagine deamination reaction. Isotopic labeling is visualized as a color gradient, with black representing 0% enrichment, red representing 50%, and purple representing 100%. **(B)** Isotopic enrichment of intracellular [I] and extracellular [E] serine and glycine following growth in media containing either uniformly labeled glucose, glycine, or serine over 6 days. The same color gradient as in (A) is applied. **(C)** Schematic representation of the aspartate transamination and asparagine deamination network before (orange) and after (dark blue) incorporation of labeled data into the mechanistic model. **(D)** Updated schematic of the serine biosynthesis pathway following incorporation of the ^13^C-labeled dataset.

Similarly, the aspartate transamination reaction which feeds into the TCA cycle via conversion of aspartate to oxaloacetate is considered an irreversible reaction in the original model (reaction 11 in **Figure 1**). However, ^13^C-isotope-labeled data for ^13^C-aspartate enrichment shows that labeled glucose, glutamate, and glutamine serve as sources of labeled aspartate (**Figure 3A**, teal-highlighted trapezoids). One plausible explanation for this labeling pattern is through a transamination reaction that converts oxaloacetate to aspartate while simultaneously converting glutamate to alpha-ketoglutarate, as illustrated in the pathway shown in **Figure 3C**. Since glucose, glutamate and glutamine feed into the TCA cycle and generate labeled TCA cycle intermediates **(Figure 1)**, which in turn produce labeled aspartate via the aspartate transamination reaction, we can reliably conclude that the aspartate transamination is indeed a reversible one as depicted in reaction 10 (**Supplementary Table 1 and Figures 1 and 3C**).

#### 3.1.2. Serine biosynthesis network

Another important modification in our model involved refining the network surrounding the amino acid serine. In the original model, the sole source of serine was interconversion from intracellular glycine and uptake of extracellular serine. However, prior research has demonstrated that serine can also be synthesized via a biosynthetic pathway originating from glycolysis, specifically through the intermediate 3-phosphoglycerate^23^. Indeed, ^13^C-isotope-labeled data indicates the accumulation of ^13^C-labeled serine coming from labeled extracellular glucose, in addition to extracellular serine and glycine. The labeling data showing the conversion of glucose to glycine and serine is shown in **Figure 3B**.

Based on this evidence, we incorporated the serine biosynthetic pathway into our updated model. In this pathway, glucose-6-phosphate is converted to the glycolytic intermediate 3-phosphoglycerate, which is subsequently transformed into 3-phosphohydroxypyruvate, then phosphoserine, and finally serine **(Figure 3D and Supplementary Figure S3)**. The full set of reactants and products across these reactions was condensed into a single stoichiometric expression, represented as Equation 35 in **Supplementary Table 1**. A corresponding kinetic expression was developed based on Michaelis-Menten kinetics. Furthermore, since incorporation of labeled extracellular serine did not yield any downstream labeling in glycolytic or TCA cycle intermediates (data not shown), we inferred that this reaction proceeds irreversibly in the current CHO model. Accordingly, Equation 35 in **Supplementary Table 1** was modeled as irreversible.

### 3.2. Model Flowchart and Computational Framework

The hybrid model consists of both kinetic equations and stoichiometric constraints. The visual flowchart illustrating the model algorithm is shown in **Figure 4**. The flowchart is divided into two main components:

1. The time course simulation (or inner loop), shown using blue arrows and shapes, which computes the concentrations of dynamically tracked metabolites, antibody, and viable cell density (VCD) over successive time points.
2. The parameter estimation loop (or outer loop), an optional process represented by red dotted arrows and solid shapes, used when parameter values need to be estimated.

**Figure 4.**
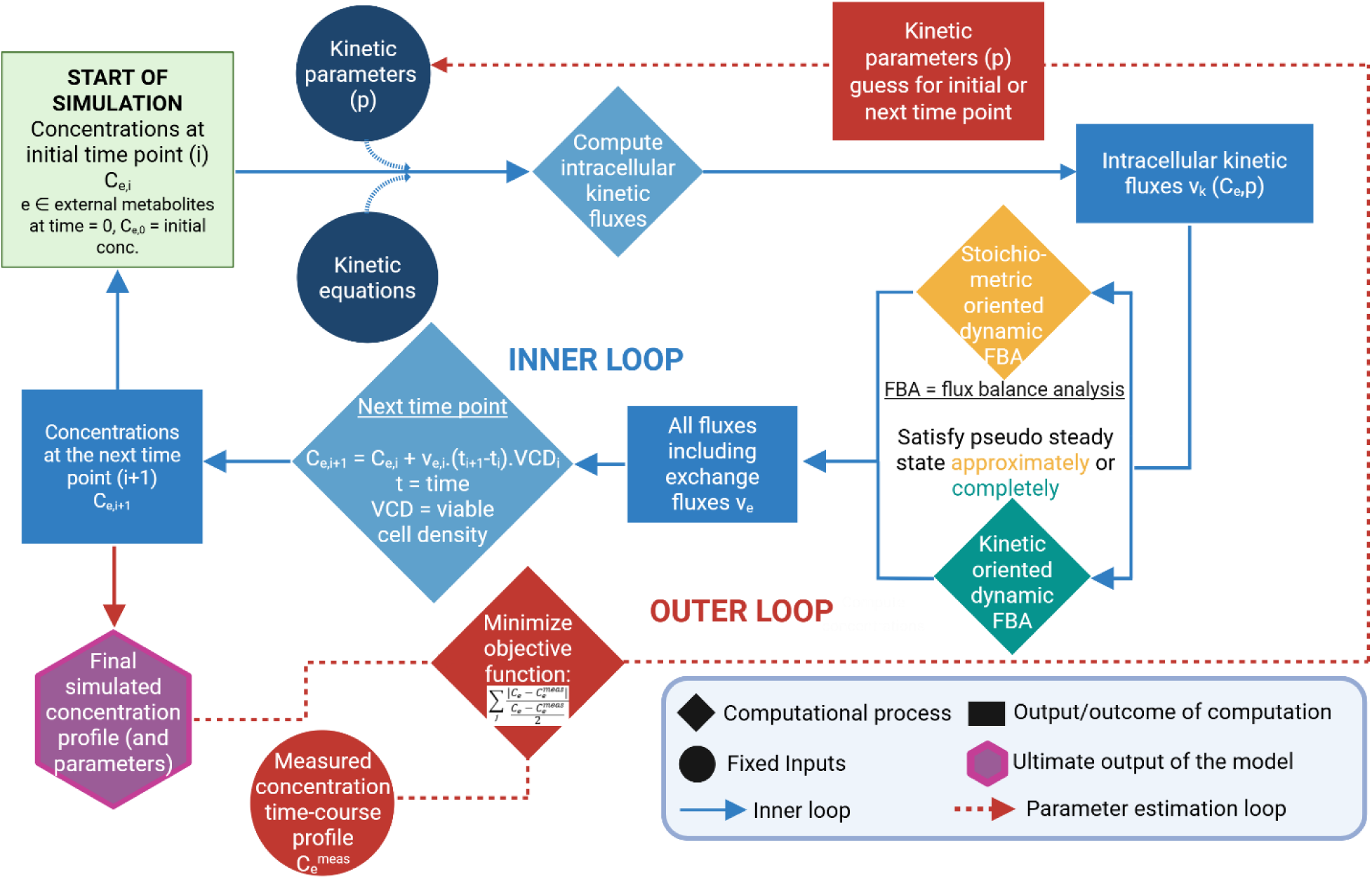
Workflow of the mechanistic model for both kinetic and stoichiometric-oriented frameworks. The model begins with initial conditions, which include extracellular metabolite concentrations, kinetic expressions, and associated parameter values. Based on these inputs, intracellular fluxes are first computed. For the stoichiometric-oriented model, the flux balance analysis (FBA) is solved completely; for the kinetic-oriented model, FBA is solved approximately. In both cases, the FBA provides values for exchange fluxes. The resulting fluxes are then used to update metabolite concentrations at the next time point using a forward Euler integration step. This process iterates until the final culture day.

The time course simulation begins with the following inputs:

1. Initial extracellular concentrations of metabolites (C_e,0_), shown in the green rectangular “Start” box
2. Fixed model inputs such as kinetic parameters (p), either as predefined values or initial guesses (if the parameter estimation loop is being run to determine the final set of parameters), shown as circles, which throughout the flowchart denote fixed inputs to the model
3. Kinetic expressions for 14 defined reactions (see **Supplementary Table 1**)

These inputs are used to compute initial intracellular fluxes for the kinetically defined reactions. This step is represented by a blue diamond, as diamonds indicate computational processes in the flowchart. The output of this step, the kinetically computed fluxes (v_k_), is depicted as a rectangle, since rectangles represent the output of a computational step. These fluxes are then used to solve the flux balance analysis (FBA), which consists of a system of stoichiometric equations based on the pseudo steady state assumption (PSSA). This assumption requires that the net flux into and out of each intracellular metabolite pool is zero, expressed as S · ν = 0, where S is the stoichiometric matrix and ν is the full set of 35 reaction fluxes.

The solving of the FBA using the pseudo steady-state assumption (PSSA) provides solutions for all fluxes. Moreover, since the system of equations in the FBA is an underdetermined system (equations (metabolites) = 24, variables (fluxes) = 35), we can have an infinite number of solutions. A subset of these fluxes is already defined by the kinetic expressions. Depending on the chosen model architecture, the FBA is solved differently:

1. Stoichiometric-oriented model-SOM **(Figure 4-yellow diamond)**: the FBA is solved in a way that selects the solution closest to the values obtained from the kinetic expressions for the 14 kinetically computed reaction fluxes (plus the exchange fluxes for biomass and antibody). This is achieved by minimizing the sum of squared differences between the fluxes computed by the kinetic equations and those determined from the FBA. As a result, the final values for all 35 fluxes, including the kinetically defined ones, are taken from the FBA solution. In this model, the pseudo steady state assumption is fully satisfied, meaning that the stoichiometric framework is prioritized while still incorporating information from kinetic expressions.

However, because this approach depends strongly on the validity of the steady state assumption and places less emphasis on empirical kinetic behavior, we also explored an alternative strategy that prioritizes the kinetic features of the model.

2. Kinetic-oriented model-KOM (**Figure 4-green diamond)**: fluxes computed from the kinetic equations are treated as final values for those specific reactions. The remaining unknown fluxes are then estimated by approximately solving the flux balance analysis. Since 14 intracellular fluxes are defined by kinetic expressions, the equation S · ν = 0 is partitioned into known and unknown components and rewritten as:

S_K._ν_K_ + S_U._ν_U_ = 0,

where, S_K_ and ν_K_ represent the submatrix and fluxes already defined by kinetic equations and S_U_ and ν_U_ represent the remaining unknowns. The introduction of fixed kinetic fluxes transforms the system into an overdetermined one, with 21 unknown fluxes and 24 metabolite balances. The ordinary least squares method is used to obtain an approximate solution. While this approach may lead to violations of the steady state condition for some metabolites, it allows the model to retain the kinetic solutions as they are, making it more suitable in cases where kinetic behavior is expected to dominate. The tradeoff is that not all intracellular fluxes will be perfectly balanced, though the model is still able to simulate realistic system behavior by preserving the empirical flux trends.

An optional parameter estimation module is used when fitting model parameters. In this case, the sum of squared differences between the full experimental dataset and the simulated time-course data is minimized.

In the flowchart:

- The **starting point** is located in the top-left corner.
- **Diamond-shaped nodes** represent computational steps.
- **Rectangles** denote inputs, outputs, or intermediate variables.
- **Circles** indicate fixed inputs.
- The **pink hexagon** represents the final model output.
- **Blue arrows** trace the inner loop that updates metabolite concentrations across time points.
- **Red arrows** represent the outer parameter estimation loop.

Figure 5A summarizes the core differences between the two models, emphasizing the expressions which are fully satisfied and those which are partially satisfied in each of the two models. Once the complete set of fluxes (ν) is determined, these values are used to update the extracellular metabolite concentrations at the next time-point (C_e,i+1_) using a forward Euler integration step, as outlined in **Supplementary Table 3**. This update step is represented by a blue diamond in the flowchart, indicating a computational process. The simulation proceeds iteratively, repeating the flux calculation and concentration update steps until the final culture day is reached. At that point, the full set of predicted concentration profiles for dynamically tracked species, antibody, and VCD is obtained. These final simulation results from the inner loop are shown in the purple hexagon (with red outline), representing the model’s final output.

**Figure 5.**
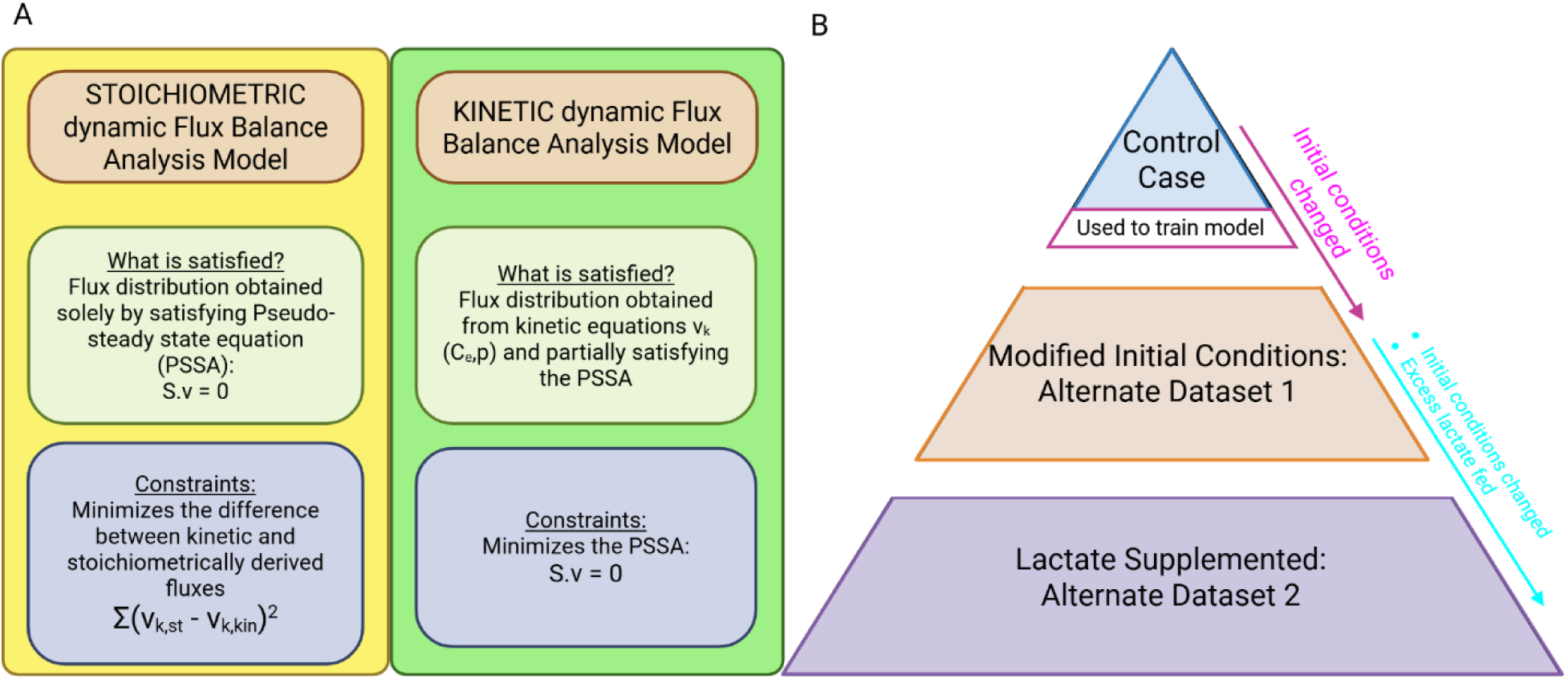
Comparison between stoichiometric-oriented (SOM) and kinetic-oriented (KOM) hybrid models, along with a description of the datasets. **(A)** Flowchart illustrating the key differences between the stoichiometric-oriented (yellow) and kinetic-oriented (green) hybrid modeling approaches. In the stoichiometric-oriented model, fluxes across the entire network are computed by fully satisfying the pseudo steady state assumption (PSSA), while minimizing the difference between the kinetically computed fluxes and those derived from the stoichiometric solution. In contrast, the kinetic-oriented model gives full preference to the kinetically computed fluxes and uses an approximate solution of the PSSA to determine the remaining fluxes in the network. **(B)** Illustration of the datasets, showing the changes from the control case (top of the pyramid) to the modified initial conditions dataset (changes in pink text), and finally to the lactate-supplemented dataset (bottom of the pyramid) (changes in blue text)

If the kinetic parameter values (p) are not predefined and require estimation, an additional input is needed: the experimentally measured concentrations and VCD across all time points. This input is represented by the red circle in the flowchart, indicating a variable that is fixed during the parameter estimation process. The model uses this experimental data to compute the objective function, shown in the red diamond, which is minimized using an optimization routine to identify the best-fitting parameter values. These parameters (red rectangle at the top of the flowchart) feed into the next iteration of the inner loop of the model. This whole process forms the outer loop of the model, illustrated by red dotted arrows, and is executed only when parameter estimation is required. In the subsequent sections, we will discuss the comparison of the model performance when the KOM is used versus the SOM for various experimental conditions.

### 3.3. Implementation of the kinetic-oriented model (KOM) for batch growth study

The KOM and SOM were implemented for a CHO cell culture producing a monoclonal antibody over a seven-day period. Cell growth (VCD), concentrations of specific amino acids, lactate levels, and IgG production were monitored throughout the culture duration. Initial conditions were based on experimental measurements (described in the methods section), and model predictions were generated for subsequent days using the approaches described previously. Kinetic parameters were fitted following the method described above in **section 2.3**, with the flowchart described in **section 3.2**. The comparison was performed for several experimental runs (Figure 5B):

1. **Control Case:** Comprises of a fixed set of initial conditions and three feeds fed at regular intervals up until the end day (day 7).
2. **Modified Initial Conditions-Alternate Dataset 1:** Similar feeding strategy as the control case but differing in the initial concentrations.
3. **Lactate-Supplementation-Alternate Dataset 2:** Differs from the control case in both initial conditions and feeding strategy and includes a bolus lactate addition to examine impact of high lactate in culture (described in the Methods and Materials section).

After running simulations using both the SOM and KOM, we evaluated their performance by comparing how well each model satisfied the pseudo-steady-state assumption (PSSA). For the SOM, the PSSA is always satisfied since one of the conditions explicitly enforced in this framework is that the PSSA must be satisfied. **Supplementary Figures S2** visually illustrates the degree to which PSSA is met for the KOM. Specifically, the heatmap shows the values of the PSSA equation over time, where green cells indicate values near zero (i.e., PSSA is satisfied) and warmer colors represent greater deviations from zero. In this model, the kinetic equations serve primarily as anchors around which the complete flux solution is determined. In contrast, the KOM does not strictly enforce the PSSA but instead minimizes deviations from it. As a result, we observed noticeable deviations for several intracellular species after a few days of simulated culture. These include alpha-ketoglutarate (AKG), ATP, FADH₂, glucose-6-phosphate (G6P), malate, NADH (cytosolic and mitochondrial), oxaloacetate (OXA), pyruvate, and oxygen. Despite these deviations, the assumption still held for many cytoplasmic metabolites in the KOM. This suggests that for dynamically tracked species, the PSSA remains reasonably valid even when it is not strictly imposed. The results indicate that the balance around these species is well defined, supporting the reliability of the KOM in capturing key metabolic dynamics.

Next, we compared the predicted concentrations of dynamically tracked species over the full duration of the culture between these two models. Figure 6 presents a side-by-side comparison of model predictions and experimental measurements for several key dynamically tracked cell culture variables. Time-course predictions for viable cell density (VCD), antibody, lactate, alanine, glutamate, aspartate, asparagine, ammonia and the serine plus glycine pool, are shown in the line graphs. The KOM (green line) placed greater emphasis on kinetic fluxes while still retaining the pseudo-steady state assumption and exhibit consistently improved agreement with experimental data (blue dots) compared to the SOM (orange line). Two of the most important parameters of biopharmaceutical cell culture processes are the viable cell culture density (VCD) and the antibody titer. Both are well-predicted by the KOM over the period of 7 days (Figure 6A **& B).** Furthermore, we also observe the correct trends for multiple other metabolites. Especially noteworthy are the predicted complex “up and down” trends for lactate and alanine starting at day 5 over the 7-day cultures (Figures 6C **and 6D).**

**Figure 6.**
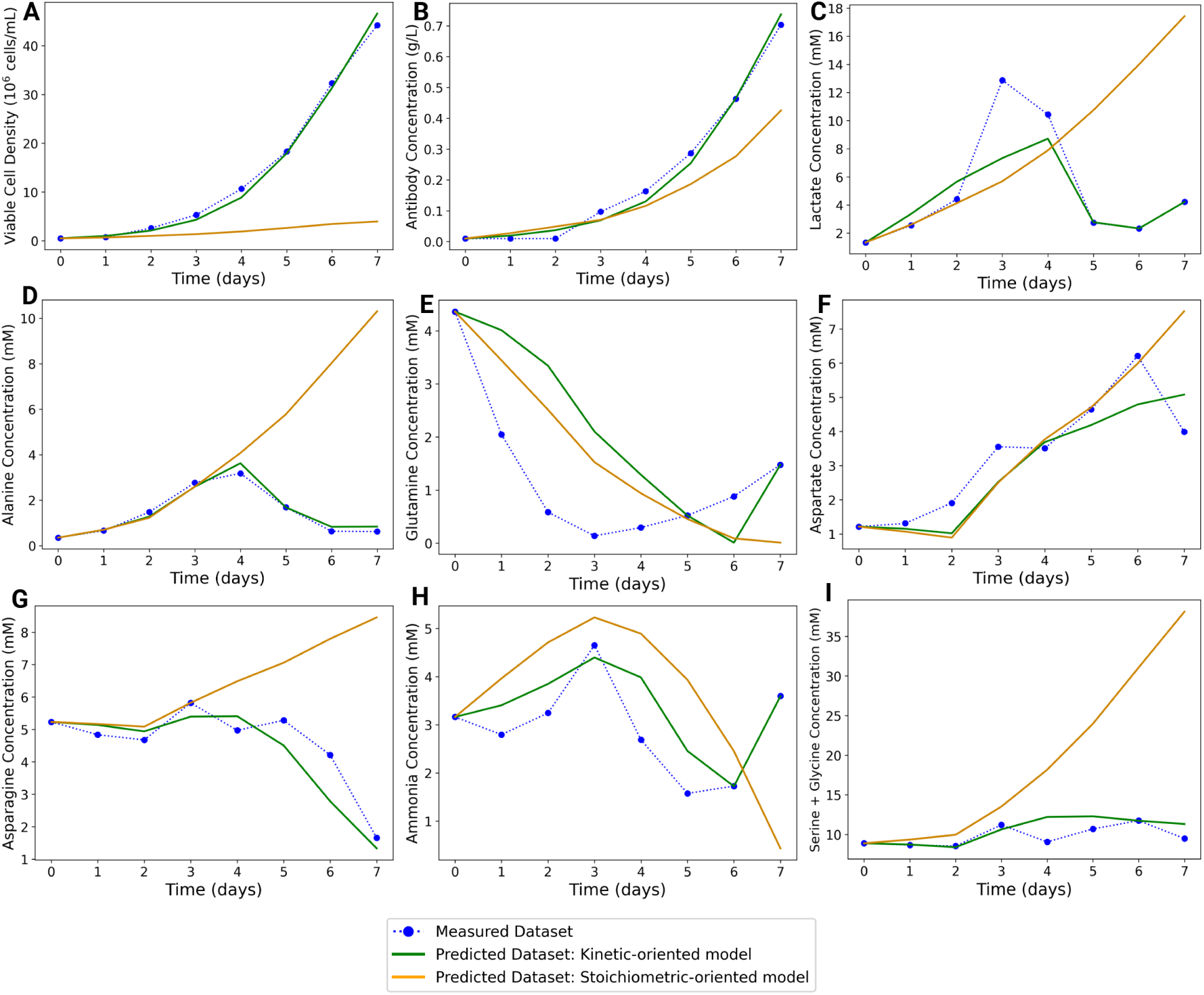
KOM vs SOM predictive profile comparison for control case. Unique parameters were obtained for the KOM and SOM, by fitting experimental data to each model. Simulation results are shown for selected dynamically tracked species from the start of culture (Day 0) to Day 7. Blue dots represent experimental measurements, green lines indicate KOM predictions, and orange lines indicate SOM predictions. Prediction profiles for **(A)** Viable Cell Density (VCD), **(B)** Antibody, **(C)** Lactate, **(D)** Alanine, **(E)** Glutamine, **(F)** Aspartate, **(G)** Asparagine, **(H)** Ammonia, **(I)** the combined Serine-Glycine pool demonstrate that the KOM largely captures observed trends, while the SOM shows lower predictive accuracy.

Importantly, the KOM successfully predicts the shift from production to consumption for both lactate and alanine using the modified kinetic expressions, without relying on factors such as the redox variable discussed earlier in **Section 2.1**. This consumption behavior is captured through the inclusion of reverse reaction terms in the kinetic equations. Specifically, the lactate equation (Reaction 2) and its associated parameters (vmax2r and Km2lac) allow for reversibility, as detailed in **Supplementary Table 1**. As lactate accumulates, the reverse term becomes increasingly dominant, leading to enhanced lactate consumption during the later stages of growth, particularly after day 4. A similar trend is observed for alanine, where consumption after day 4 is driven by the reversibility of Reaction 3, governed by parameters vmax3r and Km3ala. Since alanine and lactate are involved in the kinetic expressions of biomass and antibody synthesis respectively, this balance of lactate and alanine predictions are crucial to determining the predictions of biomass and antibody as well.

We next examine glutamine trends (Figure 6E), which are predicted reasonably well by the model. Glutamine concentration decreases sharply during the exponential phase due to its conversion to glutamate. The phenomenon of glutamine dropping during the exponential phase and accumulating at the end of the growth phase, right before the stationary phase has been shown to occur in CHO cells^20^. While the KOM underpredicts consumption of glutamine, (Figure 6E) as the predicted concentration does not fall as steeply as the measured concentration, it nonetheless captures the late-stage accumulation of glutamine, which the SOM is unable to do. Further, we observe that the trends for aspartate and asparagine (Figures 6F **and G**) are predicted reliably well. The decline in asparagine is due to its conversion to aspartate, which eventually feeds into the TCA cycle. Aspartate is seen to rise mainly due to its production from asparagine via the irreversible deamination reaction. Ammonia, involved in the aspartate-asparagine and glutamate-glutamine network is also predicted generally well (Figure 6H) with a slight initial increase followed by a decline and subsequent rise around day 7, mirroring the experimental trends.

Finally, we also observe that the serine plus glycine pool is predicted more accurately by the KOM than by the SOM (Figure 6I). Glucose was predicted more accurately by the KOM (**Figure S4A**). However, some discrepancies likely arise from the implementation of the set-point-dependent bolus glucose feed. Translating this feed from experimental conditions into the model is inherently challenging, as the abrupt fluctuations in glucose levels during the fed-batch process are difficult to reproduce in both simulations and measurements. Glutamate showed a slightly better profile with the SOM, particularly during the final two days of the simulation **(Figure S4B)**. When considering the broader context, it becomes evident that the balance of predictions of related metabolites such as ammonia and glutamine leads to an imbalance in the predictions of glutamate for the KOM. In addition, the poor predictions of cystine could be attributed to challenges in measuring cystine and its interconversion with cysteine in cell cultures.

When comparing the total error for all metabolites across all time points between the KOM and the SOM, we observe that the former has a lower overall error than the latter **(Figure S5A, first two bar graphs)**. Furthermore, **Figure S5B** presents the log-transformed ratios of errors for each individual metabolite, calculated as the error from the KOM divided by the error from the SOM. As shown in this plot, the majority of metabolites exhibit lower error values in the KOM, underscoring its improved predictive performance relative to the stoichiometric approach.

### 3.4. Implementation of Modified Initial Conditions Dataset: Alternate Dataset 1

To evaluate the robustness and generalizability of the kinetic parameters derived from the control dataset, we applied the model to another cell culture experiment with a similar feeding strategy but alternative initial conditions. The initial conditions for metabolites, amino acids, VCD and antibody varied from the control case, **Supplementary Table 4**. Once again, these initial concentrations were provided as input, and we simulated the dynamic profiles of multiple tracked species over the culture duration. The bioreactor was supplemented with three feed streams similar to the control case **(Supplementary Table 5).**

To assess the generalizability of the model, we applied the kinetic parameters optimized from the control dataset. The output variables were predicted and compared to experimental measurements (Figure 7). Once again, both viable cell density (VCD) and antibody concentrations (Figure 7A and B) are predicted with reasonable accuracy, especially when compared to the SOM. Furthermore, while the stoichiometric model fails to account for the shift in lactate dynamics from generation to consumption, the KOM is able to capture this trend, albeit with underpredictions of the levels (Figure 7C). The up and down trend for alanine is also captured well by the KOM as compared to the SOM (Figure 7D). Glutamine does not show a markedly improved profile in the KOM compared to the SOM (Figure 7E), possibly reflecting a trade-off wherein improvement in certain variables comes at the expense of others. While aspartate deviates from the measured profile at the latter stages of the cell culture, asparagine is predicted almost perfectly (Figure 7F and G). Ammonia, which is closely linked to both the aspartate-asparagine network and the glutamate-glutamine network was predicted somewhat well up to day 5 however, both models were unable to predict the rise in concentration at the latter stages (Figure 7H). The combined serine and glycine pool (Figure 7I) are all improved in the KOM but with some deviations from the measured data during the final days of culture. Finally, the glucose profile is predicted slightly better by the KOM as compared to the SOM, albeit with inaccuracies due to the reasons mentioned previously **(Figure S4C)**, and glutamate shows a generally similar trend for both the KOM and the SOM **(Figure S4D)**. Overall however, the KOM outperformed the SOM across most predicted culture variables. Once again, we plotted the total error for the kinetic-oriented and stoichiometric-oriented models **(Figure S5A, middle pair of bar graphs)**, as well as the error for each predicted variable **(Figure S5C)**. The total error was higher here than in the control, likely because the parameters were fitted to the control dataset. Nevertheless, the KOM still showed lower error than the SOM both overall and for most predicted variables. This underscores the value of prioritizing kinetics in the modeling framework, rather than relying solely on the pseudo-steady-state assumption, especially for dynamic conditions present in fed-batch CHO cell cultures.

**Figure 7.**
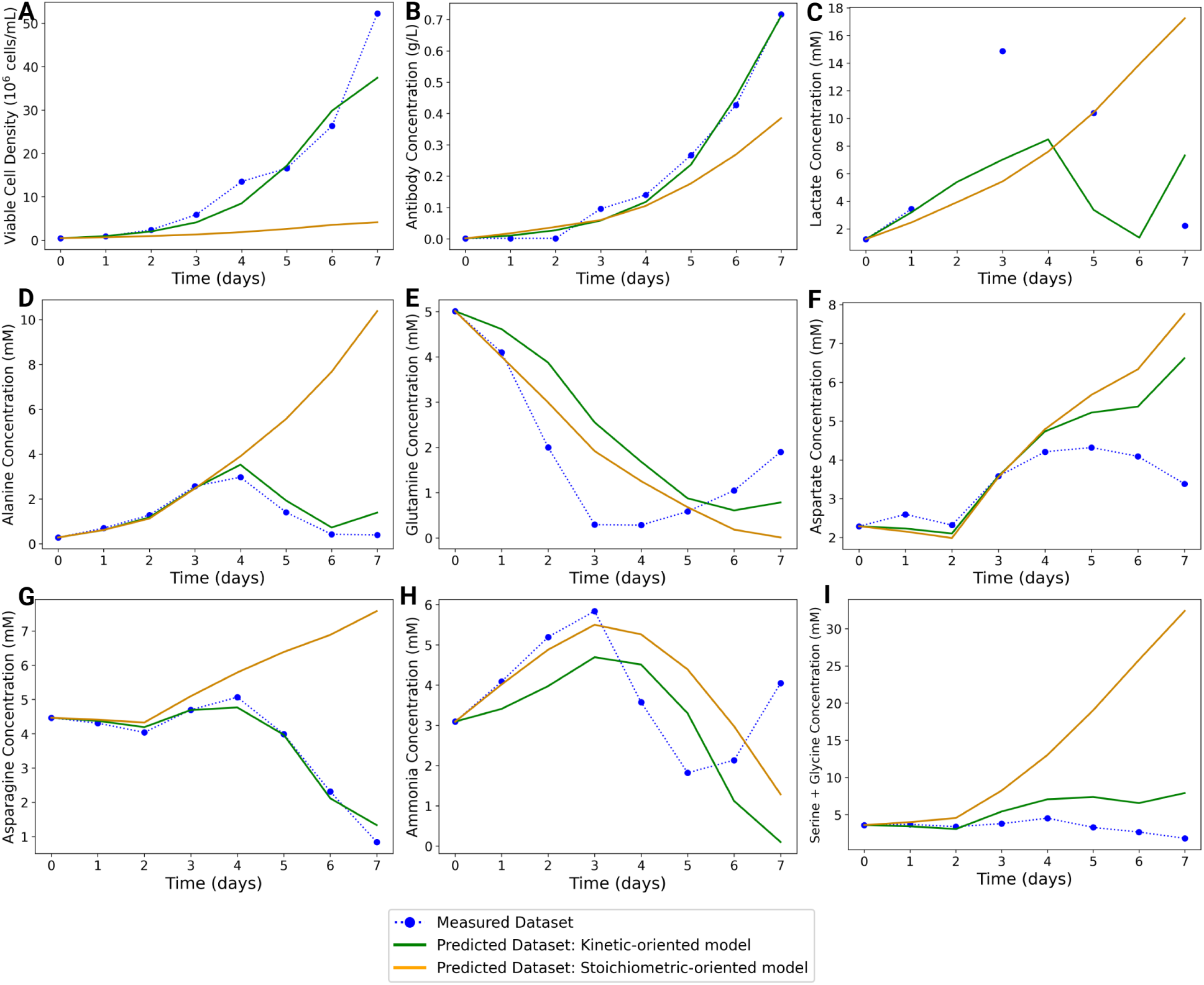
**KOM vs SOM predictive profile comparison for Modified Initial Conditions-Alternate Dataset 1**. Parameters used for prediction were those obtained by from the control case fit. Simulation results are shown for selected dynamically tracked species from the start of culture (Day 0) to Day 7. Blue dots represent experimental measurements, green lines indicate KOM predictions, and orange lines indicate SOM predictions. Prediction profiles for **(A)** Viable Cell Density (VCD), **(B)** Antibody, **(C)** Lactate, **(D)** Alanine, **(E)** Glutamine, **(F)** Aspartate, **(G)** Asparagine, **(H)** Ammonia, **(I)** the combined Serine-Glycine pool demonstrate that the KOM largely captures observed trends, while the SOM shows lower predictive accuracy, similar to the observations made in the control case dataset.

We also compared the model fits and errors with parameters that were derived from directly fitting to the Modified Initial Conditions dataset. Fitting this dataset produced final simulation errors that were slightly lower than those obtained with the control case parameters for both model types (**Figure S6A)**. The SOM did not provide good fits, consistent with previous observations for this framework (data not shown).

Furthermore, the error for the SOM, as shown in **Figure S6A**, remained much higher than that of the KOM for both sets of simulations, i.e., using the control case parameters, and the newly fit parameters. In addition, we compared the predictions obtained using the control case parameters against those obtained with the fit parameters for the KOM **(Figure S6B-E)**. While the VCD predictions were largely similar between the two parameters sets **(Figure S6B)**, some other dynamically tracked metabolites and amino acids were predicted better with the newly fit parameters. This included glutamine, for which the new parameters captured both the decline and subsequent rise in concentration, unlike the control-case parameters **(Figure S6C)**. The aspartate profile was also better predicted with the new parameters, reflecting its consumption during the latter stages of the culture, as compared to the control-case parameters **(Figure S6D)**. Ammonia, a key metabolite closely linked to both of these amino acids also showed improved alignment with experimental data **(Figure S6E)**.

Overall, the KOM with parameters fitted to the Modified Initial Conditions dataset achieved slightly better agreement with experimental profiles than the simulations based on the more generalizable control-case parameters, and the total error was only slightly reduced for the newly fit parameters in both the kinetic and the stoichiometric-oriented models **(Figure S6A)**, suggesting that the initial parameters can in general be effective and retained across different runs, at least under similar process and feed conditions.

### 3.5. Implementation of Lactate-Supplemented Case: Alternate Dataset 2

To further assess the performance of the KOM and the parameters derived from the control dataset, we applied the model to a third cell culture experiment which differs from the control case in both initial conditions **(Supplementary Table 4 and** Figure 5B) as well as in feeding inputs (as described in the Methods and Materials section). As before, the initial concentrations of substrates and key metabolites were specified as input **(Supplementary Table 4)**, and the dynamic profiles of multiple tracked species were simulated over the culture period.

Shown in Figure 8 are the results of the lactate-supplemented case in which some of the key process parameters were evaluated. Viable cell density (VCD) was very poorly predicted by the SOM, consistent with its performance across all three datasets, while the KOM overpredicted VCD (Figure 8A). Antibody production was also noticeably overpredicted by the KOM compared to the experimental data, whereas the stoichiometric model showed closer agreement (Figure 8B). Interestingly the KOM was able to predict the late switch to production for the case with control parameters. In contrast, the SOM failed to adequately capture the lactate consumption phase that followed lactate production and supplementation, while the KOM reproduced the lactate profile reasonably well over the 7-day culture period after accounting for the added lactate (Figure 8C). It is important to note that the kinetic parameters used in this simulation were originally fitted to the control case, where the peak lactate concentration was 11.54 mM. Under the excess lactate condition, however, measured lactate levels reached 38.2 mM, likely exceeding the valid range for those parameters. This mismatch is reflected in the inaccurate predictions of VCD and IgG and aligns with the known inhibitory effects of elevated lactate on both cell growth and antibody production ^24^.

**Figure 8.**
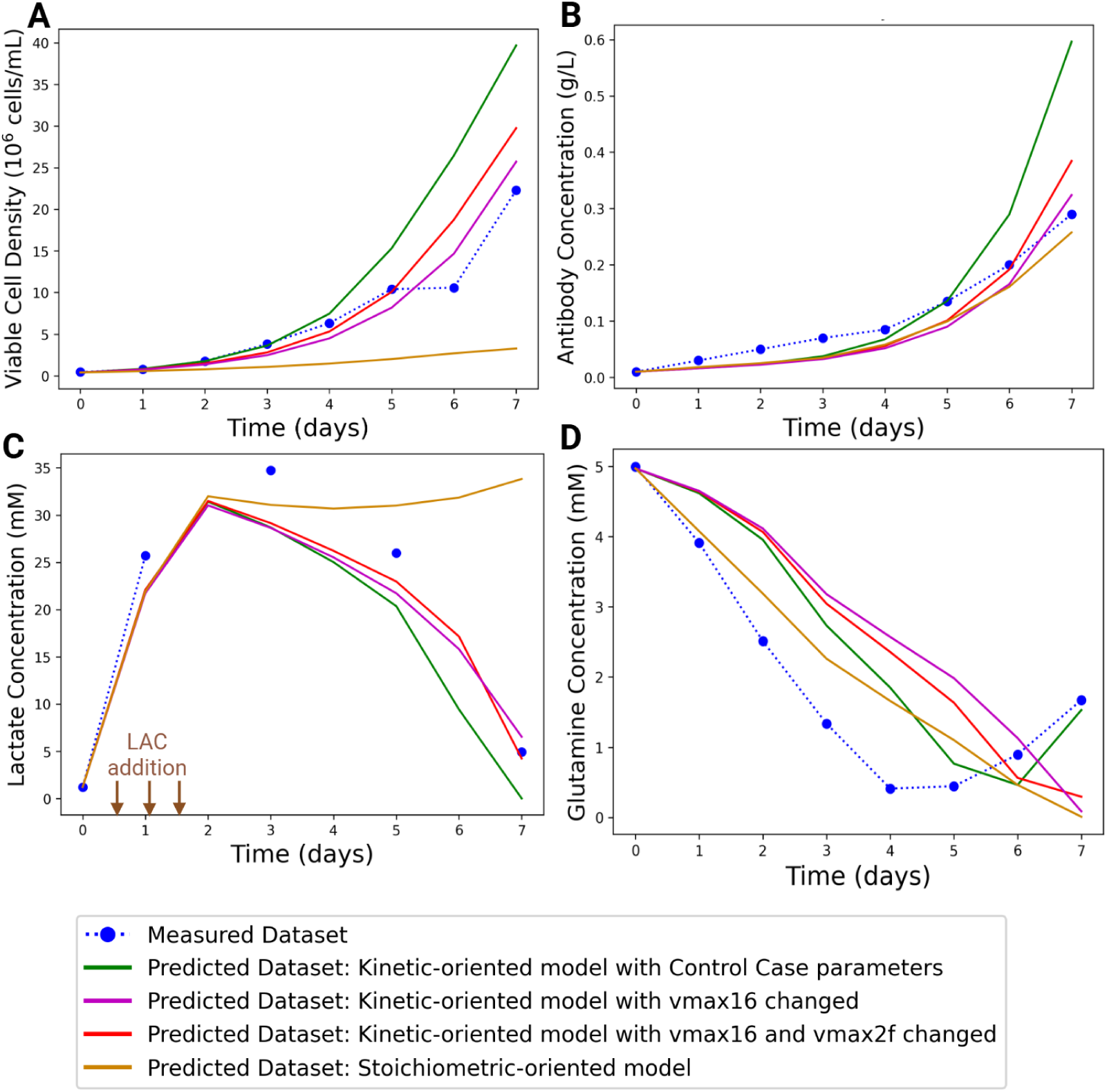
KOM vs SOM predictive profile comparison for Lactate-Supplemented Case-Alternate Dataset 2. Lactate additions (days 0.5, 1, and 1.5) are indicated by brown arrows in panel C. Simulation results are shown for selected dynamically tracked species from Day 0 to Day 7. Blue dots represent experimental measurements, green lines indicate KOM predictions using control-case parameters, and orange lines indicate SOM predictions. Additional fits were obtained by reducing kinetic parameter values: vmax16 (VCD-related, red line) and vmax2f (Lactate-related, purple line). Profiles are shown for **(A)** Viable Cell Density (VCD), **(B)** Antibody, **(C)** Lactate, **(D)** Glutamine

Furthermore, lactate explicitly inhibits antibody production through the kinetic expression for antibody (Reaction 17 in **Supplementary Table 1).** Therefore, the slight underprediction of lactate by the KOM likely contributed, at least in part, to the overprediction of the antibody profile under the current parameter set.

From the results for VCD across all three datasets, we observed that the SOM predicts only a very modest increase in VCD. To investigate the cause of this severe underperformance, we examined the flux profiles of biomass generation (which directly contributes to VCD) and of associated amino acids involved in the kinetic expression for biomass generation, particularly alanine, whose profile is poorly predicted by the stoichiometric model. In **Supplementary** Figure 7 **and Supplementary Table 7**, the flux profiles for v16 (biomass generation) and v22 (alanine generation/consumption) are shown for both the kinetic-and stoichiometric-oriented models for the control case. For the KOM, v16 (green line) and v22 (blue line) display strong fluctuations, which is consistent with the feeding regime that drives large changes in metabolite concentrations during culture. These variations are appropriately captured by the kinetic model. By contrast, the stoichiometric model shows relatively flat profiles for fluxes of biomass (v16, yellow line) and alanine (v22, red line). The flux for biomass generation is much lower than expected, leading to an underprediction of VCD. At the same time, the flux for alanine remains high and consistently positive, causing the alanine concentration to continue rising rather than falling in the later stages of culture, as observed in the experimental data. Given that alanine plays an important role in the biomass generation expression, the biomass and by extension, VCD is poorly predicted by the SOM. Taken together, these results strongly suggest that both the magnitudes and trends of key fluxes were not adequately captured by the SOM. In other words, this framework was unable to represent the dynamic behavior of the CHO fed-batch system.

Next, we compared the total error across this dataset, which revealed that the cumulative error across all metabolites and time points is consistently higher for the SOM than for the KOM **(Figure S5A, last pair of bar graphs)**. The total error for this lactate supplemented dataset was lower as compared to other datasets, primarily due to the smaller number of measured species available for evaluation. The log-transformed ratios of errors for each individual metabolite also shows that the majority of metabolites across all datasets exhibit lower error values in the KOM **(Figure S5D)**, highlighting its improved predictive performance relative to the stoichiometric approach for all the three datasets.

To address these deficiencies in the kinetic-oriented model’s capacity to describe the effects of high lactate, we allowed specific kinetic parameters associated with biomass production and lactate generation to vary and evaluated the predictions for VCD, IgG, lactate, and several other major metabolites measured in this follow-up simulation. Specifically, we reduced the value of vmax16, the rate constant for the biomass reaction (red line in Figure 8) and also decreased vmax2f, the forward rate constant associated with lactate generation, in another simulation (magenta line in Figure 8). These changes resulted in a lower predicted biomass formation rate and, consequently, a reduction in VCD values that were more in line with experimental measurements. Antibody predictions also improved under this updated parameter set (Figure 8B, red and magenta). Meanwhile, the KOM retained its capacity to predict the expected trend of lactate production followed by consumption. Furthermore, the fit with experimental lactate data was improved on days 6 and 7 (Figure 8C). Glutamine (Figure 8D) showed the same underprediction of consumption observed in previous cases, possibly due to the influence of ammonia. Finally, glucose predictions were also improved for the KOM compared to the SOM **(Figure S4E)** as was the trend for glutamate **(Figure S4F)**. Overall, these results indicate that the modified KOM, using parameters originally derived from the control dataset, is superior to the SOM. However, parameter adjustments were useful to apply the model to experimental conditions such as lactate supplementation that differ significantly from those used during the initial parameterization.

## 4. Conclusion

Accurately simulating CHO cell metabolism is useful for optimizing cell culture media design and enhancing bioprocess performance. Our study demonstrates that incorporating ^13^C-labeled data into a hybrid model that combines FBA and kinetic modeling provides a principled approach to model construction. This integration leads to improved predictive accuracy of CHO cell metabolism across a range of growth conditions. The modifications to the reaction stoichiometries based on ^13^C-labeled data, including updates to asparagine and aspartate reactions and the addition of a serine biosynthesis network, enabled refinement of the model. The hybrid model dynamically tracked metabolite concentrations while iteratively calculating flux distributions, offering significant advantages over using FBA or kinetic modeling alone. Simplifying changes, such as removing the redox variable (R) and temperature-sensitive parameters, further streamlined the model. These adjustments not only accelerated parameter estimation but also reduced unnecessary complexity, making the model more adaptable to a variety of metabolomics datasets involving glucose, lactate, and amino acids.

Ultimately, these refinements reduced total error, producing predictions more closely aligned with experimental observations.

One key finding of this study was the superior performance of kinetic-oriented simulations compared to those prioritizing stoichiometric balances. The KOM framework allows for time-dependent tracking of metabolite concentrations without strictly enforcing the pseudo-steady-state assumption, instead relying on empirical kinetic expressions.

Including realistic terms, such as the reverse reaction components for rate expressions of lactate and alanine production, facilitates the metabolic shift of these key cellular metabolites and enables more accurate predictions of their production-to-consumption shifts. The SOM, by contrast, may require additional parameters such as the redox variable included in the prior model^12^, to explicitly capture the metabolic shift from lactate production to consumption. The KOM approach can capture this behavior without such add-ons, which in turn allows it to better capacity to reproduce dynamic phenomena such as metabolite accumulation, consumption shifts, and transient behaviors that are often observed in CHO fed-batch cultures. In contrast, the SOM is limited by its steady-state constraints, which can hinder accurate simulation of key outputs such as viable cell density, antibody titer, and amino acid profiles over extended culture periods.

Furthermore, our results highlight the enhanced predictive capabilities of the hybrid model in CHO fed-batch systems under various conditions such as modified baseline conditions and lactate supplementation case. We generated parameters from the control case and successfully applied it to the alternate datasets, producing predictions that are largely consistent with experimental data. This was particularly evident for the dataset with altered initial conditions, where the generalizable parameters performed nearly as well as the fitted parameters, with only minor differences. The model reliably predicts key metabolites, viable cell culture density (VCD), and antibody production, supporting its practical utility in improving CHO cell culture performance. In the lactate supplemented case, the critical trend of lactate production followed by consumption was still captured. Additional parameter adjustments further improved predictions for lactate as well as VCD and antibody levels. While discrepancies remain for some metabolites, the overall performance demonstrates that emphasizing kinetic expressions while partially satisfying FBA constraints yields more accurate and generalizable results.

Future enhancements could focus on integrating additional metabolic pathways such as the pentose phosphate pathway (PPP), hexosamine biosynthesis, and even fatty acid metabolism branching from carbohydrate pathways^25,26^. While more detailed models are available, these are primarily stoichiometric in nature to avoid the large number of kinetic parameters required for kinetic or hybrid models^25,27,28^. Incorporating black-or gray-box models to represent relationships not captured with current models could supplement the hybrid framework and improve its predictive capacity^29^, factoring in additional variables such as pH and osmolarity that may also influence antibody production, VCD, and other cell culture parameters. Most importantly, the hybrid modeling approach with a kinetic oriented focus described here represents a valuable foundational approach for building robust and practical tools to enhance CHO cell culture bioproduction, ultimately contributing to more efficient and scalable biomanufacturing processes.

## Acknowledgements

This study was funded by the Advanced Mammalian Biomanufacturing Innovation Center (AMBIC) through the Industry-University Cooperative Research Center Program under U.S. National Science Foundation grant numbers 1624684 and 2100800.

Experimental work was supported by the grant numbers IIP-162464 and EEC-2100442 and was partially supported by NSF grant number OIA-1736123 The funder played no role in study design, data collection, analysis and interpretation of data, or the writing of this manuscript.

## Contributions

P.K, N.N.: Writing – original draft, Validation, Investigation, Formal analysis, Data curation, Software; S.K.: Writing – original draft, Data collection, Formal analysis, Data curation; E.M., T.B.: Investigation, Formal analysis, Software; Y.K.: Writing – review, Software, Supervision, Conceptualization; S.H.: Writing – review, Data curation, Supervision, Funding acquisition, Conceptualization; M.J.B.: Writing – review & editing, Visualization, Supervision, Project administration, Funding acquisition, Data curation, Conceptualization.

## Data availability

All data generated or analyzed during this study are included in this published article and its supplementary information files.

## Code availability

The underlying code for this study is not publicly available but may be made available to qualified researchers on reasonable requests from the corresponding author.

## References

1. Zhu, M. M., Mollet, M., Hubert, R. S., Kyung, Y. S. & Zhang, G. G. Industrial Production of Therapeutic Proteins: Cell Lines, Cell Culture, and Purification. in Handbook of Industrial Chemistry and Biotechnology (eds. Kent, J. A., Bommaraju, T. V & Barnicki, S. D.) 1639–1669 (Springer International Publishing, Cham, 2017). doi:10.1007/978-3-319-52287-6_29.

2. Glaser, V. Meeting Current Needs and Assessing Future Opportunities to Drive the Global Bioeconomy. Industrial Biotechnology 9, 271–274 (2013).

3. Lu, R. M. et al. Development of therapeutic antibodies for the treatment of diseases. J Biomed Sci 27, (2020).

4. Dumont, J., Euwart, D., Mei, B., Estes, S. & Kshirsagar, R. Human cell lines for biopharmaceutical manufacturing: history, status, and future perspectives. Critical Reviews in Biotechnology vol. 36 1110–1122 (2016).

5. Okamura, K., Badr, S., Murakami, S. & Sugiyama, H. Hybrid Modeling of CHO Cell Cultivation in Monoclonal Antibody Production with an Impurity Generation Module. Ind Eng Chem Res 61, 14898–14909 (2022).

6. Almquist, J., Cvijovic, M., Hatzimanikatis, V., Nielsen, J. & Jirstrand, M. Kinetic models in industrial biotechnology - Improving cell factory performance. Metab Eng 24, 38–60 (2014).

7. Sha, S., Huang, Z., Wang, Z. & Yoon, S. Mechanistic modeling and applications for CHO cell culture development and production. Curr Opin Chem Eng 22, 54–61 (2018).

8. Stalidzans, E., Seiman, A., Peebo, K., Komasilovs, V. & Pentjuss, A. Model-based metabolism design: Constraints for kinetic and stoichiometric models. Biochem Soc Trans 46, 261–267 (2018).

9. Regueira, A., Bevilacqua, R., Mauricio-Iglesias, M., Carballa, M. & Lema, J. M. Kinetic and stoichiometric model for the computer-aided design of protein fermentation into volatile fatty acids. Chemical Engineering Journal 406, (2021).

10. Øyås, O. & Stelling, J. Genome-scale metabolic networks in time and space. Curr Opin Syst Biol 8, 51–58 (2018).

11. Ben Yahia, B., Malphettes, L. & Heinzle, E. Macroscopic modeling of mammalian cell growth and metabolism. Appl Microbiol Biotechnol 99, 7009–7024 (2015).

12. Nolan, R. P. & Lee, K. Dynamic model of CHO cell metabolism. Metab Eng 13, 108–124 (2011).

13. Kuriya, Y. & Araki, M. Dynamic flux balance analysis to evaluate the strain production performance on shikimic acid production in Escherichia coli. Metabolites 10, (2020).

14. Mahadevan, R., Edwards, J. S. & Doyle, F. J. Dynamic Flux Balance Analysis of diauxic growth in Escherichia coli. Biophys J 83, 1331–1340 (2002).

15. MathWorks. fmincon (Version R2023a). https://www.mathworks.com/help/optim/ug/fmincon.html (2023).

16. Chitwood, D. G. et al. Microevolutionary dynamics of eccDNA in Chinese hamster ovary cells grown in fed-batch cultures under control and lactate-stressed conditions. Sci Rep 13, (2023).

17. Harcum, S. W. et al. PID controls: the forgotten bioprocess parameters. Discover Chemical Engineering 2, (2022).

18. Synoground, B. F. et al. Transient ammonia stress on Chinese hamster ovary (CHO) cells yield alterations to alanine metabolism and IgG glycosylation profiles. Biotechnol J 16, (2021).

19. Naik, H. M. et al. Elucidating uptake and metabolic fate of dipeptides in CHO cell cultures using 13C labeling experiments and kinetic modeling. Metab Eng 83, 12–23 (2024).

20. Zhang, L. xiang et al. Responses of CHO-DHFR cells to ratio of asparagine to glutamine in feed media: cell growth, antibody production, metabolic waste, glutamate, and energy metabolism. Bioresour Bioprocess 3, (2016).

21. Xu, P., Dai, X. P., Graf, E., Martel, R. & Russell, R. Effects of glutamine and asparagine on recombinant antibody production using CHO-GS cell lines. Biotechnol Prog 30, 1457– 1468 (2014).

22. Kirsch, B. J. et al. Metabolic analysis of the asparagine and glutamine dynamics in an industrial Chinese hamster ovary fed-batch process. Biotechnol Bioeng 119, 807–819 (2022).

23. Kim, H. & Park, Y. J. Links between Serine Biosynthesis Pathway and Epigenetics in Cancer Metabolism. Clin Nutr Res 7, 153 (2018).

24. Lao, M. S. & Toth, D. Effects of ammonium and lactate on growth and metabolism of a recombinant Chinese hamster ovary cell culture. Biotechnol Prog 13, 688–691 (1997).

25. Martínez, J. A., Bulté, D. B., Contreras, M. A., Palomares, L. A. & Ramírez, O. T. Dynamic Modeling of CHO Cell Metabolism Using the Hybrid Cybernetic Approach With a Novel Elementary Mode Analysis Strategy. Front Bioeng Biotechnol 8, (2020).

26. Robitaille, J., Chen, J. & Jolicoeur, M. A single dynamic metabolic model can describe mAb producing CHO cell batch and fed-batch cultures on different culture media. PLoS One 10, (2015).

27. Pan, X., Dalm, C., Wijffels, R. H. & Martens, D. E. Metabolic characterization of a CHO cell size increase phase in fed-batch cultures. Appl Microbiol Biotechnol 101, 8101–8113 (2017).

28. Jiménez del Val, I., et al. CHOmpact: A reduced metabolic model of Chinese hamster ovary cells with enhanced interpretability. Biotechnol Bioeng 120, 2479–2493 (2023).

29. Cui, T. et al. Data-driven and physics informed modeling of Chinese Hamster Ovary cell bioreactors. Comput Chem Eng 183, (2024).

