## Supplementary Information for "Comparing Kinetic versus Stoichiometric Priorities in Hybrid Models of CHO Metabolism"

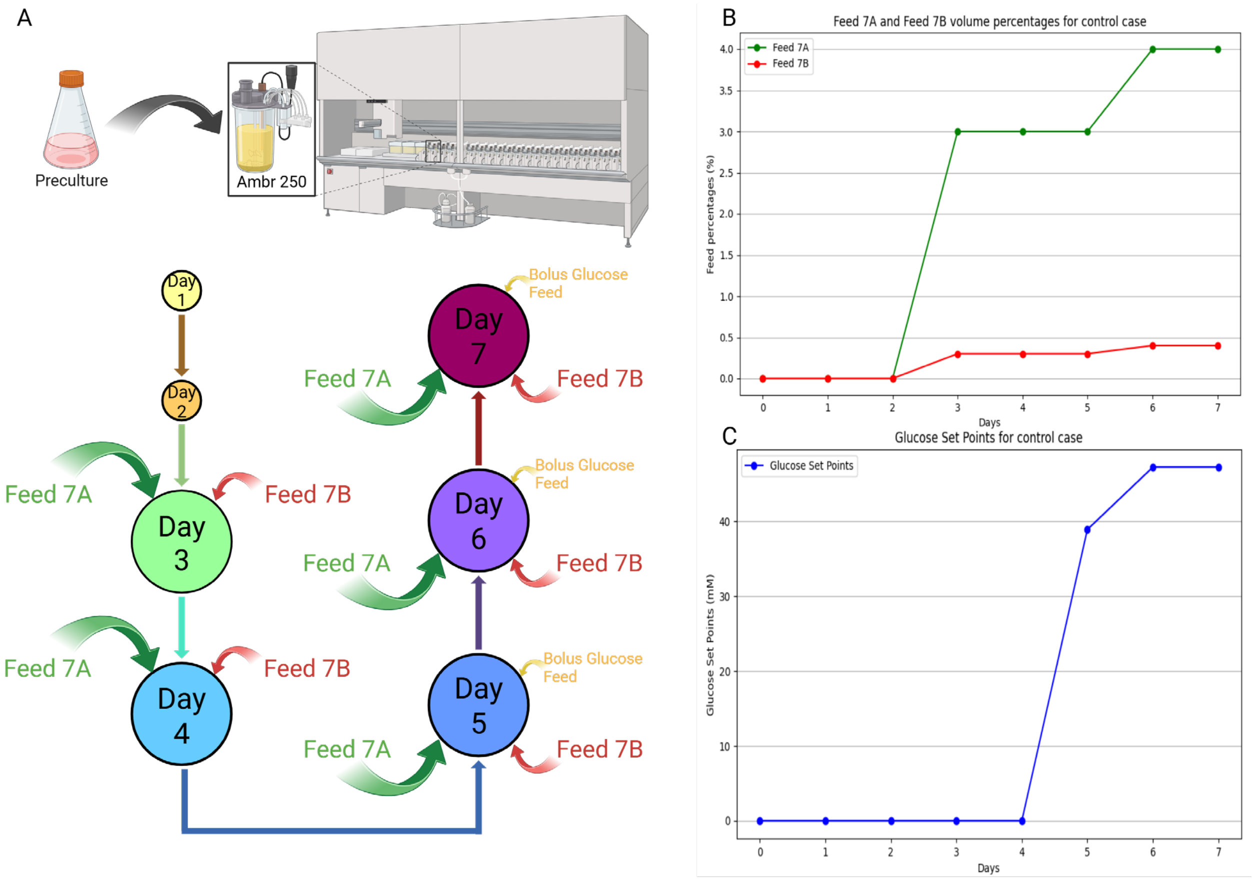


**Supplemental Figure 1: Schematic of feeding strategy, feed concentrations, and glucose set points. (A)** A schematic representation of the CHO cell culture from Day 0 to Day 7, highlighting the feeding strategy. Cells were precultured in shake flasks and subsequently transferred to an Ambr250 vessel. Feeding begins on Day 3 and includes Feed 7A, Feed 7B (Cell Boost 7a/7b- Cytiva), and bolus glucose additions. **(B)** Corresponding line graphs for Feed 7A and 7B concentrations. **(C)** Glucose set points.

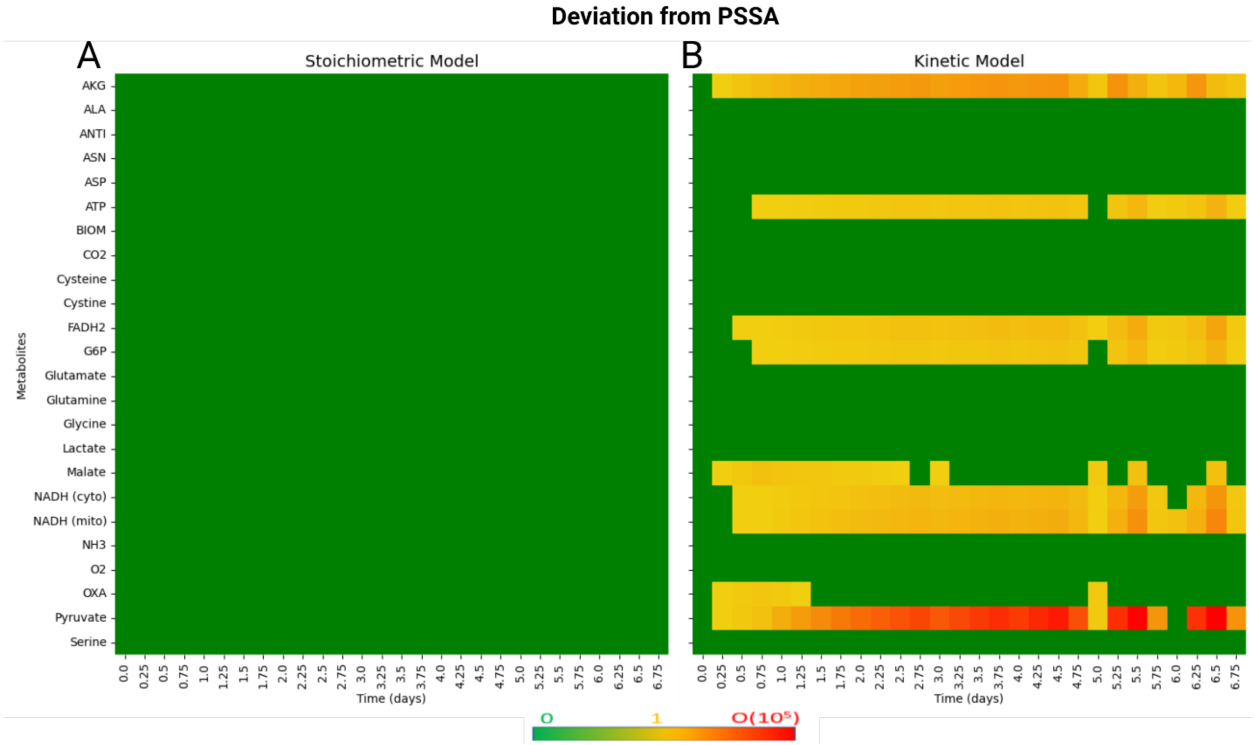


**Supplemental Figure 2: Comparison of PSSA performance in stoichiometric-oriented vs. kinetic-oriented models.** At each simulated time point, the S·v matrix is solved to evaluate the extent to which intracellular fluxes satisfy the pseudo steady state assumption (PSSA). **(A)** In the stoichiometric-oriented model, the fluxes are computed such that the PSSA is strictly enforced at each time point. The resulting heatmap shows that the net flux across each metabolite in the control dataset remains near zero, indicating full adherence to steady state conditions. **(B)** In the kinetic-oriented model, the fluxes from the kinetic expressions are fixed inputs, and therefore the PSSA cannot be fully satisfied. As a result, several metabolites show moderate to significant deviations from PSSA. Specifically, AKG, ATP, FADH2, G6P, malate, and oxaloacetate exhibit moderate deviations, while NADH and pyruvate, both in the cytosol and mitochondria, show substantial deviations.

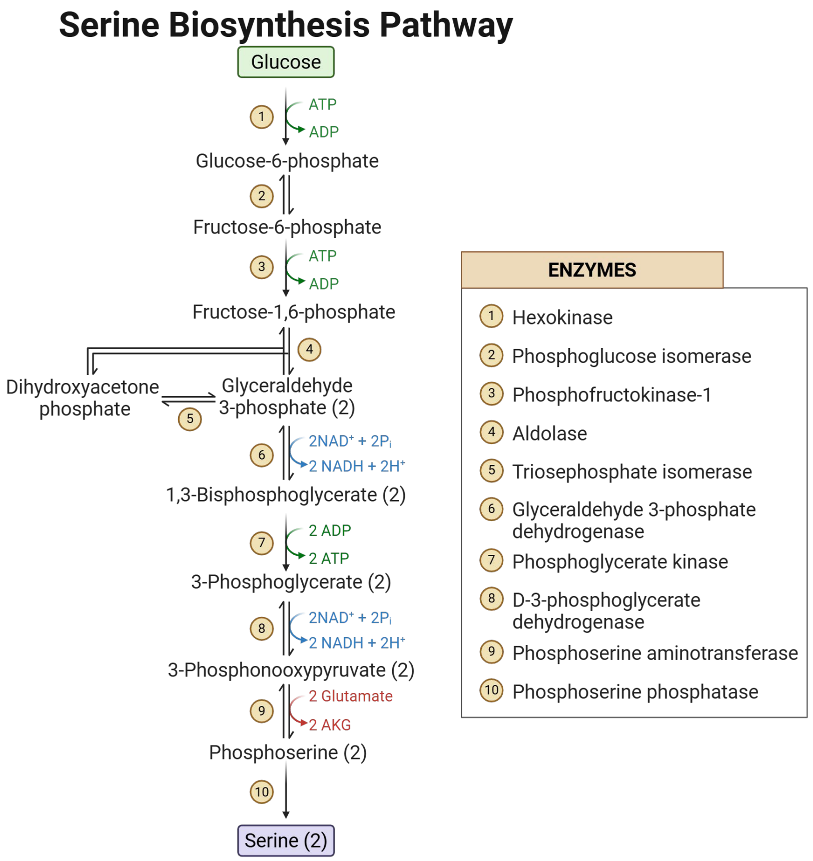


**Supplemental Figure 3.** Individual reactions in the serine biosynthesis pathway, from which a condensed reaction expression was derived.

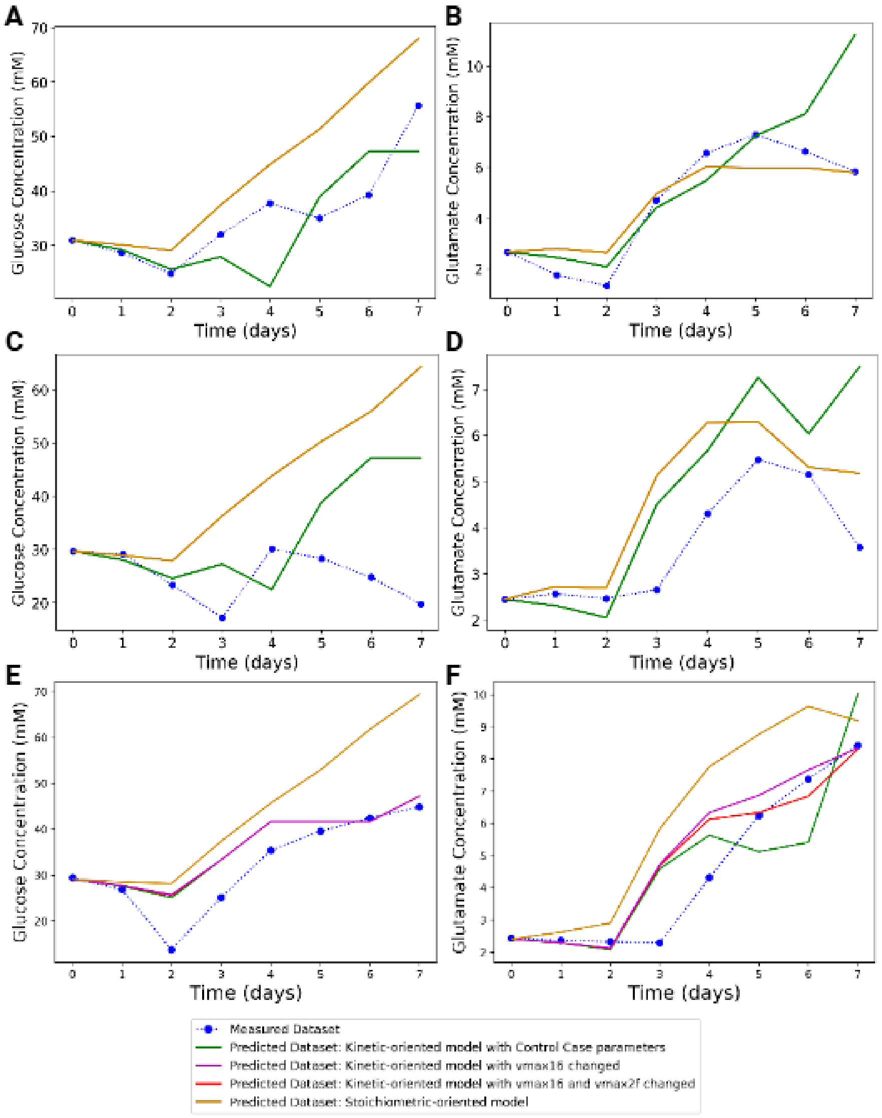


**Supplemental Figure 4. KOM vs SOM predictive profile comparison for glucose (left side graphs) and glutamate (right side graphs) profiles for (A-B)** Control case (Parameters were fit to this dataset) **(C-D)** Modified Initial Conditions- Alternate Dataset 1 (control case parameters used) **(E-F)** Lactate-Supplemented Case- Alternate Dataset 2 (control case parameters used, with modifications to vmax16 and vmax2f parameters)

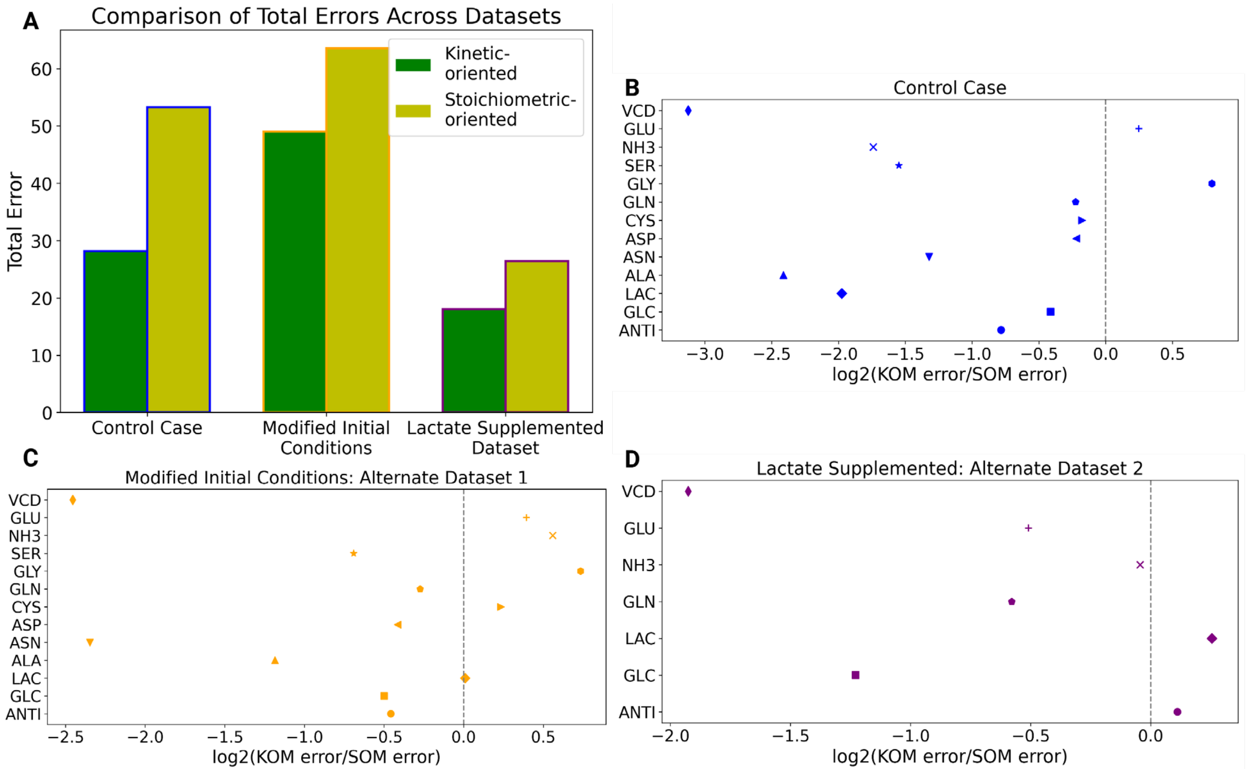


**Supplementary Figure 5: Analysis of error between the kinetic-oriented model (KOM) and the stoichiometric-oriented model (SOM). (A)** Total error for the KOM versus the SOM across all datasets, computed as the sum of errors for all metabolites and timepoints. **(B)** Error comparison for the Control Case, shown as log2 fold change of KOM error relative to SOM error for each metabolite. Most metabolites fall below zero, indicating lower error for the KOM. **(C)** Error comparison for the Modified Initial Conditions (Alternate Dataset 1), shown as log2 fold change of KOM error relative to SOM error. The majority of metabolites fall below zero, again favoring the KOM. **(D)** Error comparison for the Lactate-Supplemented Case (Alternate Dataset 2), shown as log2 fold change of KOM error relative to SOM error, shown only for those metabolites with measured data available. Most metabolites fall below zero, indicating reduced error with the KOM.

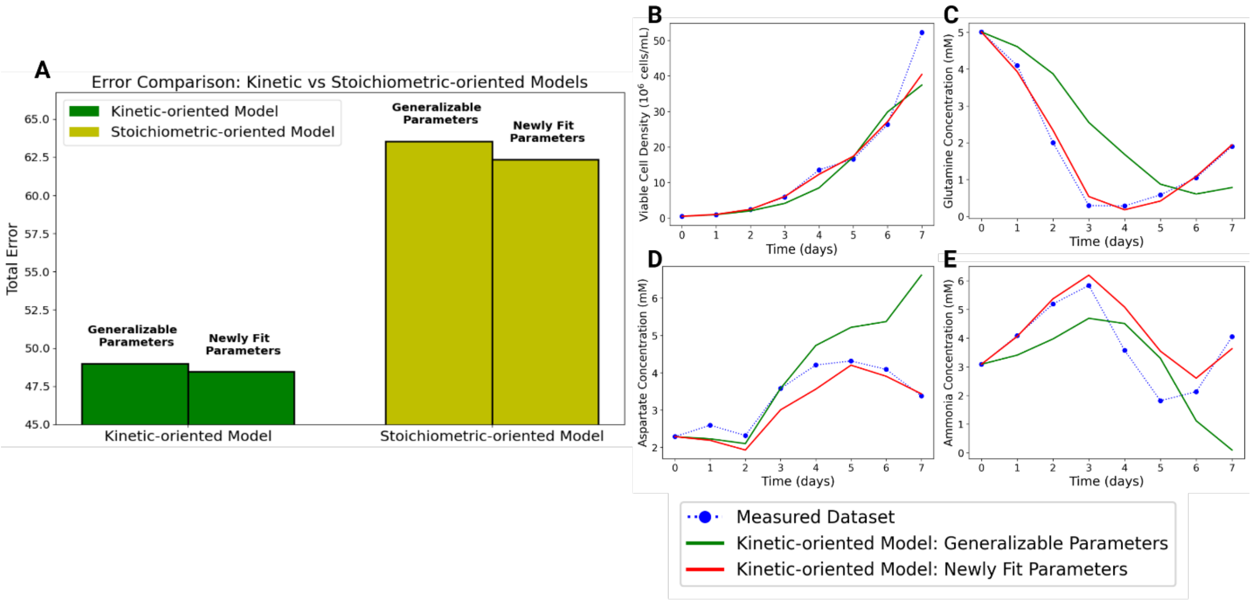


**Supplementary Figure 6: Comparison of model fits and errors using parameters derived from the Modified Initial Conditions dataset. (A)** Total simulation error for KOM and SOM with parameters from either the Control Case or the Modified Initial Conditions dataset, showing consistently lower error for KOM. **(B–E)** KOM predictions using control-case versus newly fit parameters for selected profiles: **(B)** Viable Cell Density (VCD), **(C)** Glutamine, **(D)** Aspartate, and **(E)** Ammonia. Newly fit parameters improved predictions for Glutamine, Aspartate, and Ammonia, while VCD remained similar between parameter sets.

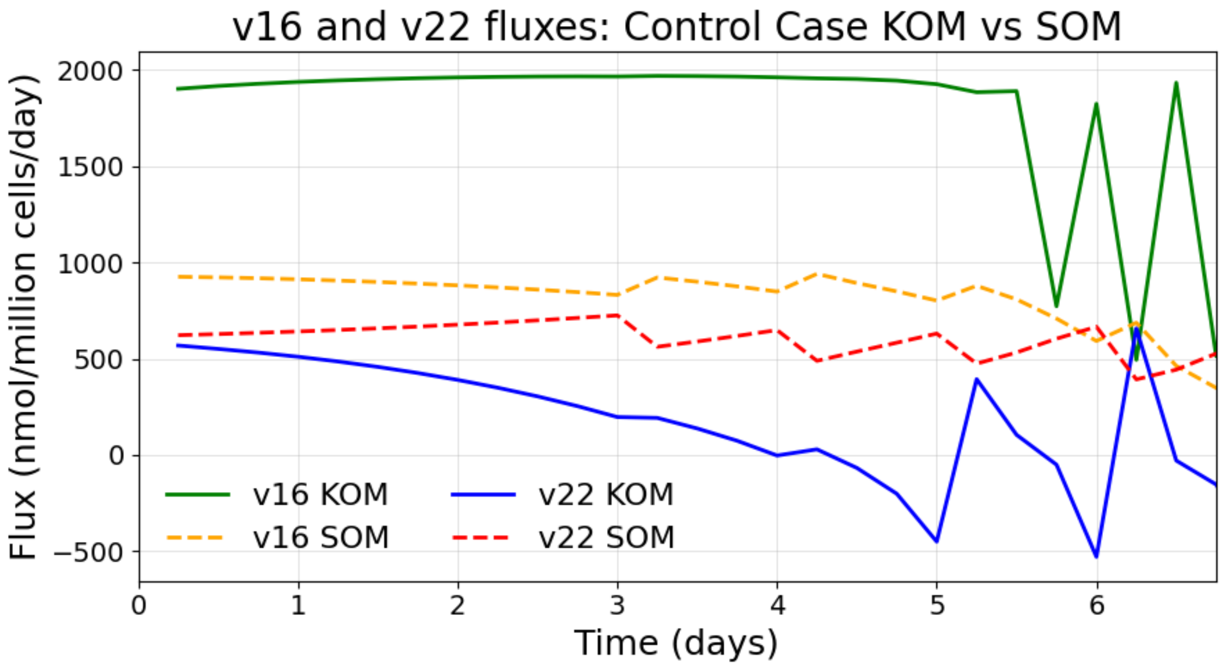


**Supplementary Figure 7: Flux profiles for biomass generation (v16) and alanine generation/consumption (v22) in the Control Case.** The KOM captures dynamic fluctuations in biomass (v16, green) and alanine (v22, blue) fluxes consistent with feeding-driven metabolite changes, while the SOM produces flat profiles for biomass (v16, yellow) and alanine (v22, red). Reduced biomass flux leads to underprediction of VCD, and persistently positive alanine flux results in continued accumulation, highlighting the inability of the SOM to capture dynamic culture behavior.

**Table 1:** Kinetic expressions for 14 reactions, including 12 intracellular reactions and 2 exchange fluxes, with their corresponding reaction indices. All expressions follow the Michaelis-Menten framework and are defined by extracellular concentrations of dynamically tracked variables together with kinetic parameters. The kinetic parameters include maximum reaction velocity ($v_{max}$), half-saturation constant ($K_m$), and inhibition constant ($K_i$).

| Index number | Reaction | Kinetic expression |
|---|---|---|
| R1 | G6P→2PYR+3ATP+ 2NADH | $v_1 = \frac{v_{max1} \cdot \left(\frac{GLC}{K_{m1}}\right)}{\left(1 + \frac{LAC}{K_{i1}}\right) \cdot \left(1 + \frac{GLC}{K_{m1}}\right)}$ |
| R2 | PYR + NADH → LAC | $v_2 = \frac{v_{max2f} \cdot \left[\left(\frac{GLC}{K_{m2glc}}\right)\right] - v_{max2r} \cdot \left(\frac{LAC}{K_{m2lac}}\right)}{1 + \left(\frac{LAC}{K_{m2lac}}\right) + \left(\frac{GLC}{K_{m2glc}}\right)}$ |
| R3 | PYR+GLU→ALA+ AKG | $v_3 = \frac{v_{max3f} \cdot \left[\left(\frac{GLC}{K_{m3glc}}\right)\right] - v_{max3r} \cdot \left(\frac{ALA}{K_{m3ala}}\right)}{1 + \left(\frac{GLC}{K_{m3glc}}\right) + \left(\frac{ALA}{K_{m3ala}}\right)}$ |
| R8 | GLN→GLU+NH3 | $v_8 = \frac{v_{max8f} \cdot \left(\frac{GLN}{K_{m8gln}}\right) - v_{max8r} \cdot \left(\frac{GLU}{K_{m8glu}}\right)\left(\frac{NH3}{K_{m8nh3}}\right)}{1 + \left(\frac{GLN}{K_{m8gln}}\right) + \left(\frac{GLU}{K_{m8glu}}\right) + \left(\frac{NH3}{K_{m8nh3}}\right) + \left(\frac{GLU}{K_{m8glu}}\right)\left(\frac{NH3}{K_{m8n}}\right.}$ [illegible] |
| R9 | AKG+NH3+NADH→ GLU | $v_9 = \frac{v_{max9f} \cdot \left(\frac{NH3}{K_{m9nh3}}\right) - v_{max9r} \cdot \left[\left(\frac{GLU}{K_{m9glu}}\right) + \left(\frac{GLN}{K_{m9gln}}\right)\right]}{1 + \frac{NH3}{K_{m9nh3}} + \frac{GLU}{K_{m9glu}} + \frac{GLN}{K_{m9gln}} + \left(\frac{GLU}{K_{m9glu}}\right)\left(\frac{GLN}{K_{m9gln}}\right)}$ |
| R10 | ASN→ASP+NH3 | $v_{10} = \frac{vmax_{10f} \cdot \left(\frac{ASN}{K_{m10asn}}\right)}{1 + \left(\frac{ASN}{K_{m10asn}}\right)}$ |
| R11 | ASP+AKG→OXA+ GLU | $v_{11} = \frac{v_{max11f} \cdot \left[\left(\frac{ASP}{K_{m11asp}}\right) + \left(\frac{ASN}{K_{m11asn}}\right)\right] - v_{max11r}\left(\frac{GLU}{K_{m11glu}}\right)}{1 + \left(\frac{ASP}{K_{m11asp}}\right) + \left(\frac{ASN}{K_{m11asn}}\right) + \left(\frac{ASP}{K_{m11asp}}\right)\left(\frac{ASN}{K_{m11asn}}\right)}$ |
| R12 | SER+CO2+NH3+NADH→2 GLY | $v_{12} = \frac{v_{max12f} \cdot \left(\frac{SER}{K_{m12ser}}\right)\left(\frac{NH3}{K_{m12nh3}}\right) - v_{max12r} \cdot \left(\frac{GLY}{K_{m12gly}}\right)^2}{1 + \left(\frac{SER}{K_{m12ser}}\right) + \left(\frac{NH3}{K_{m12nh3}}\right) + \left(\frac{GLY}{K_{m12gly}}\right) + \left(\frac{GLY}{K_{m12gly}}\right)^2}$ |
| R13 | Cystine+NADH→ 2Cysteine | $v_{13} = \frac{v_{max13} \cdot \left(\frac{C-C}{K_{m13}}\right)}{\left(1 + \left(\frac{C-C}{K_{m13}}\right)\right)}$ |
| R16 | Biomass production | $v_{16} = \frac{v_{max16} \cdot \left(\frac{GLN}{K_{m16a}}\right) \cdot \left(\frac{ASN}{K_{m16b}}\right) \cdot \left(\frac{ALA}{K_{m16c}}\right)}{1 + \left(\frac{GLN}{K_{m16a}}\right) + \left(\frac{ASN}{K_{m16b}}\right) + \left(\frac{ALA}{K_{m16c}}\right) + \left(\frac{GLN}{K_{m16a}}\right) \cdot \left(\frac{ASN}{K_{m16b}}\right) + \left(\frac{GLN}{K_{m16a}}\right) \cdot \left(\frac{ALA}{K_{m16c}}\right) + \left(\frac{ASN}{K_{m16b}}\right) \cdot \left(\frac{ALA}{K_{m16c}}\right) + \left(\frac{GLN}{K_{m16a}}\right) \cdot \left(\frac{ASN}{K_{m16b}}\right) \cdot \left(\frac{ALA}{K_{m16c}}\right)}$ |
| R17 | Antibody synthesis | $v_{17} = \frac{v_{max17}}{1 + \left(\frac{LAC}{K_{i17}}\right)}$ |

| | | |
|---|---|---|
| R18 | Biomass exchange reaction | $v_{18} = v_{16}$ |
| R19 | Antibody exchange reaction | $v_{19} = v_{17}$ |
| R35 | G6P + ATP + 2GLU→ 2NADH + 2SER + 2AKG | $v_{35} = \frac{v_{max35} \cdot \left(\frac{GLC}{K_{m35a}}\right) \cdot \left(\frac{GLU}{K_{m35b}}\right)^2}{\left(1 + \left(\frac{GLC}{K_{m35}}\right) + \left(\frac{GLU}{K_{m35b}}\right) + \left(\frac{GLU}{K_{m35b}}\right)^2\right)}$ |

**Table 2:** Reaction expressions for all 35 reactions in the model. Extracellular metabolites are denoted with the subscript “E”. NADH is separated into cytosolic and mitochondrial pools, represented by the subscripts “cyto” and “mito”, respectively.

| | |
|---|---|
| 1. | G6P → 2 PYR + 3 ATP + 2 $NADH_{cyto}$ |
| 2. | PYR + $NADH_{cyto}$ ↔ LAC |
| 3. | PYR + GLU ↔ ALA + AKG |
| 4. | PYR + OXA → AKG + 2 $CO_2$ + 2 $NADH_{mito}$ |
| 5. | AKG → MAL + CO2 + $NADH_{mito}$ + FADH2 + ATP |
| 6. | MAL → OXA + $NADH_{mito}$ |
| 7. | MAL → PYR + $CO_2$ |
| 8. | GLN ↔ GLU + $NH_3$ |
| 9. | AKG + $NH_3$ + $NADH_{mito}$ ↔ GLU |
| 10. | ASN → ASP + $NH_3$ |
| 11. | ASP + AKG ↔ OXA + GLU |
| 12. | SER + $CO_2$ + $NH_3$ + $NADH_{cyto}$ ↔ 2 GLY |
| 13. | Cystine + $NADH_{cyto}$ → 2 Cysteine |
| 14. | $NADH_{mito}$ + 0.5 $O_2$ → 2.5 ATP |
| 15. | $FADH_2$ + 0.5 $O_2$ → 1.5 ATP |
| 16. | 0.0838 ALA + 0.041 ASN + 0.0804 ASP + 8.6825 ATP + 0.0261 Cysteine + 0.452 G6P + 0.0873 GLN + 0.056 GLY + 0.427 OXA + 0.096 SER → BIOM + 0.004 $FADH_2$ + 0.0082 GLU + 0.4445 MAL + 0.6391 $NADH_{mito}$ + 0.2085 PYR |
| 17. | 0.0614 ALA + 0.0344 ASN + 0.0389 ASP + 9.2 ATP + 0.024 Cysteine + 0.0479 GLU + 0.0449 GLN + 0.0719 GLY + 0.1 SER → ANTI |
| 18. | BIOM → $BIOM_E$ |
| 19. | ANTI → $ANTI_E$ |
| 20. | $GLC_E$ + ATP → G6P |
| 21. | LAC ↔ $LAC_E$ |
| 22. | ALA ↔ $ALA_E$ |
| 23. | $ASN_E$ → ASN |
| 24. | ASP ↔ $ASP_E$ |
| 25. | $Cystine_E$ + GLU → Cystine + $GLU_E$ |
| 26. | $GLN_E$ ↔ GLN |
| 27. | GLY ↔ $GLY_E$ |
| 28. | $SER_E$ ↔ SER |
| 29. | NH3 ↔ $NH3_E$ |
| 30. | $O2_E$ → O2 |
| 31. | CO2 → $CO2_E$ |
| 32. | 2 Cysteine + O2 → Cystine |
| 33. | GLU ↔ $GLU_E$ |
| 34. | $NADH_{cyto}$ → 0.5 $NADH_{mito}$ + 0.5 $FADH_2$ |
| 35. | G6P + ATP + 2 GLU → 2 $NADH_{cyto}$ + 2 SER + 2 AKG |

**Table 3:** Mass balances for all dynamically tracked species were modeled using forward Euler ODEs. Overall cell density was calculated from biomass, while a separate ODE described dead cell density.

| **Numerical Integration and Biomass Conversion Formulas** |
| --- |
| Forward Euler equations for metabolite species: $$C_{k,i} = C_{k,i-1} + \frac{v_{k,i-1}(t_i - t_{i-1})VCD_{i-1}}{1000}$$ C = Concentration (mM)<br>k = metabolite species or Biomass or Antibody<br>VCD = viable cell density (million cells/mL)<br>v = exchange flux (nmol/million cells/day)<br>$t_i$ = $i^{th}$ timepoint (days) |
| Overall cell density calculated as: $$Overall\ cell\ density = \frac{Biomass}{2.31}$$ Since it was experimentally determined that 1 million cells is equivalent to 2.31mM of biomass Nolan et al.[12]. |
| Dead cell density calculated as: $$Dead\ cell\ density_i = Dead\ cell\ density_{i-1} + k_d VCD_{i-1}$$ $k_d$ = Death rate constant |

**<u>Table 4:</u>** Measured initial concentrations of metabolites, amino acids, antibody, and cell densities for each of the three experiments: control case, modified initial conditions case, and lactate-supplemented case.

| **Measured Initial Concentrations/Quantities of Metabolites** | | | |
|---|---|---|---|
| **Metabolite/species** | **Control Case** | **Modified Initial Conditions: Alternate Dataset 1** | **Lactate-Supplemented: Alternate Dataset 2** |
| Antibody (g/L) | 0.079 | 0.01 | 0.079 |
| Glucose (mM) | 30.9 ± 0.2 | 29.7 ± 0.9 | 29.1 ± 0.4 |
| Lactate (mM) | 1.33 | 1.26 ± 0.04 | 1.22 |
| Alanine (mM) | 0.35 ± 0.14 | 0.28 ± 0.02 | N/A |
| Asparagine (mM) | 5.23 ± 0.22 | 4.47 ± 0.11 | N/A |
| Aspartate (mM) | 1.21 ± 0.18 | 2.29 ± 0.05 | N/A |
| Cystine (mM) | 0.332 ± 0.092 | 1.49 ± 0.54 | N/A |
| Glutamine (mM) | 4.36 ± 0.01 | 5.01 ± 0.13 | 4.995 ± 0.025 |
| Glycine (mM) | 1.12 ± 0.03 | 0.609 ± 0.424 | N/A |
| Serine (mM) | 7.76 ± 0.72 | 2.98 ± 2.69 | N/A |
| Ammonia (mM) | 3.16 ± 0.02 | 3.09 ± 0.02 | 3.09 |
| Glutamate (mM) | 2.67 ± 0.01 | 2.45 ± 0.075 | 2.4 |
| Viable cell density- VCD (million cells/mL) | 0.495 ± 0.005 | 0.452 ± 0.015 | 0.395 ± 0.054 |

Values are presented as mean ± SEM when n ≥ 2; for timepoints with a single replicate, only the individual value is shown. In the lactate-supplemented condition, some amino acids were not measured and are indicated as N/A.

**Table 5:** Concentrations of select model-relevant metabolites in each of the three feed streams.

| **Concentrations/Quantities (mM) Of Metabolites in Feeds** | | | |
|---|---|---|---|
| **Metabolites** | **Feed 7A** | **Feed 7B** | **Bolus Glucose** |
| Glucose | 360.8 | | 2500 |
| Alanine | 21.8 | | |
| Asparagine | 35 | | |
| Aspartate | 62.3 | | |
| Cysteine/Cystine | 0 | 85.00 | |
| Glutamine | 0 | | |
| Glycine | 0 | | |
| Serine | 110.5 | | |
| Ammonia | 0 | | |
| Glutamate | 100.6 | | |

**Table 6**: PID settings for DO control in the ambr250 HT bioreactor system. Level 0 represents the initial conditions for the control manipulated variables. Level 1 increases the total gas flow rate when the DO falls below the DO set-point. No overlay gas was used. Air, $O_2$, and $CO_2$ are all provided through the sparge gas stream.

| Level | Variable | Minimum | Maximum | $K_p$ | $t_I$ (s) | $t_D$ (s) |
|---|---|---|---|---|---|---|
| 0 | Stir speed (rpm)<br>Gas flow rate (mL/min)<br>$O_2$ mix (%) *<br>$CO_2$ flow rate (mL/min) | 300<br>2<br>0<br>0.1 | 300<br>2<br>0<br>0.1 | NA | NA | NA |
| 1 | Total gas flow rate (mL/min) | 2 | 20 | 0.10 | 0 | 100 |
| 2 | $O_2$ mix (%) | 0 | 50 | 0.15 | 0 | 100 |
| 3 | Stir speed (rpm) | 300 | 400 | 2.0 | 0 | 100 |
| 4 | $O_2$ mix (%) | 50 | 100 | 0.15 | 0 | 250 |
| 5 | Stir speed (rpm) | 400 | 800 | 1.0 | 0 | 100 |
| 6 | $O_2$ flow (mL/min) | 0 | 20 | 0.50 | 0 | 100 |

*$O_2$ mix is the percentage of oxygen mixed with the sparge gas, which contains air and a variable amount of carbon dioxide that is used for pH control. NA – Not Applicable

**Table 7:**

| | Fluxes for biomass generation and alanine production/consumption | | | |
|---|---|---|---|---|
| Timepoint (days) | Kinetic-oriented model | | Stoichiometric-oriented model | |
| | v16 (Biomass) | v22 (Alanine) | v16 (Biomass) | v22 (Alanine) |
| 0.25 | 1902.75881 | 568.766402 | 926.1325 | 622.4039 |
| 0.5 | 1917.54014 | 551.1775735 | 922.8625 | 628.4039 |
| 0.75 | 1929.10597 | 532.1731374 | 918.4553 | 634.7456 |
| 1 | 1938.68794 | 510.511173 | 912.9556 | 641.7611 |
| 1.25 | 1946.57271 | 485.8383579 | 906.461 | 649.4838 |
| 1.5 | 1952.98885 | 457.7788081 | 899.0158 | 657.9429 |
| 1.75 | 1958.11387 | 425.9740235 | 890.6229 | 667.1657 |
| 2 | 1962.07746 | 390.0703879 | 881.2488 | 677.1773 |
| 2.25 | 1964.95975 | 349.7056774 | 870.8234 | 687.9992 |
| 2.5 | 1966.78315 | 304.4755899 | 859.2348 | 699.6447 |
| 2.75 | 1967.49319 | 253.8617468 | 846.3195 | 712.1111 |
| 3 | 1966.91826 | 197.088443 | 831.8425 | 725.3644 |
| 3.25 | 1969.9147 | 192.5624972 | 922.343 | 562.1384 |
| 3.5 | 1968.90104 | 137.6963393 | 900.3383 | 589.5616 |
| 3.75 | 1966.66259 | 73.80820885 | 876.1293 | 618.5277 |
| 4 | 1962.44631 | -2.87612554 | 849.1936 | 648.8931 |
| 4.25 | 1957.40174 | 29.12342583 | 941.0681 | 489.1707 |
| 4.5 | 1953.90091 | -67.4800315 | 893.2149 | 537.202 |
| 4.75 | 1945.5732 | -201.808235 | 850.1037 | 583.3925 |
| 5 | 1926.94057 | -450.552445 | 802.2661 | 630.6819 |
| 5.25 | 1885.26551 | 393.9662387 | 879.255 | 474.5714 |
| 5.5 | 1890.93178 | 104.5555484 | 808.4232 | 532.5969 |
| 5.75 | 772.913055 | -50.5703094 | 708.887 | 604.2199 |
| 6 | 1825.10499 | -529.401199 | 592.3594 | 666.2646 |
| 6.25 | 494.299326 | 657.5740704 | 687.3929 | 392.5595 |
| 6.5 | 1934.52876 | -29.2845556 | 463.0169 | 443.3808 |
| 6.75 | 513.463542 | -153.695668 | 348.5713 | 526.9391 |
| 7 | 1783.68125 | -528.63431 | 285.5604 | 606.7188 |
